# Integrated analysis of anatomical and electrophysiological human intracranial data

**DOI:** 10.1101/230912

**Authors:** Arjen Stolk, Sandon M. Griffin, Roemer van der Meij, Callum Dewar, Ignacio Saez, Jack J. Lin, Giovanni Piantoni, Jan-Mathijs Schoffelen, Robert T. Knight, Robert Oostenveld

## Abstract

The exquisite spatiotemporal precision of human intracranial EEG recordings (iEEG) permits characterizing neural processing with a level of detail that is inaccessible to scalp-EEG, MEG, or fMRI. However, the same qualities that make iEEG an exceptionally powerful tool also present unique challenges. Until now, the fusion of anatomical data (MRI and CT images) with the electrophysiological data and its subsequent analysis has relied on technologically and conceptually challenging combinations of software. Here, we describe a comprehensive protocol that addresses the complexities associated with human iEEG, providing complete transparency and flexibility in the evolution of raw data into illustrative representations. The protocol is directly integrated with an open source toolbox for electrophysiological data analysis (FieldTrip). This allows iEEG researchers to build on a continuously growing body of scriptable and reproducible analysis methods that, over the past decade, have been developed and employed by a large research community. We demonstrate the protocol for an example complex iEEG data set to provide an intuitive and rapid approach to dealing with both neuroanatomical information and large electrophysiological data sets. We explain how the protocol can be largely automated, taking under an hour to complete, and readily adjusted to iEEG data sets with other characteristics.

## INTRODUCTION

Intracranial EEG (iEEG) allows simultaneous recordings from tens to hundreds of electrodes placed directly on the neocortex (electrocorticography, ECoG), or intracortically (stereoelectroencephalography, SEEG). In humans, the most common implementation of iEEG is when non-invasive techniques such as scalp-EEG and MRI do not provide sufficient information to guide surgery in medication refractory epilepsy patients. Each electrode reflects the activity of tens of thousands of neurons^1, 2^, and the recording and stimulation of these neural populations allow for identification of epileptogenic zones, as well as for mapping of functionally eloquent areas of human cortex to guide neurosurgery. The outcome of these procedures can be directly observed when the neural or behavioral response is straightforward such as speech arrest or muscle movement with direct stimulation^3^. Any more complex empirical study requires accurate knowledge of an electrode’s location in relation to the brain’s anatomy that is linked to the local electrophysiological signal. This integrated information is key to basic and clinical research work aimed at understanding human neural and cognitive processing^4, 5^.

Human iEEG analysis has traditionally relied on stand-alone and ad hoc workflows for the separate analysis of anatomical and functional aspects of the iEEG data, presenting researchers with a series of challenges to realize the full potential of this exceptionally powerful tool. To process the neuroanatomical data research labs are tasked with assembling software combinations for the conversion of file formats (e.g., DICOM to NIfTI using MRIConvert), coregistration of anatomical scans (e.g., CT to MRI using SPM^6^, FSL^7^, or AFNI^8^), localization of electrodes (e.g., BioImage Suite^9^), and the sorting and labeling of electrodes to match the format of the functional recording file (manually, or using custom software). This technological obstacle is receiving increasing attention in the form of more efficient workflows for localizing and visualizing electrodes^10–16^, but no protocol exists that allows researchers to efficiently process the anatomical data within a single work environment, and seamlessly fuse with the electrophysiological data and its subsequent analysis. Ideally, in the light of scientific reproducibility^17^, such a protocol should also provide complete transparency in the evolution of raw data into results and illustrative representations, allowing for a convenient and efficient exchange of data and workflows between researchers. These two components are particularly valuable in a growing field where the analysis of data is uniquely complex, but where the gold standard for that analysis is yet to be defined.

Here, we describe - at the implementation level - a comprehensive protocol to address the series of challenges associated with both the anatomical and functional aspects of human iEEG analysis. The protocol is directly integrated with the MATLAB-based open source FieldTrip toolbox (Box 1), offering the opportunity to readily and flexibly build on a continuously growing set of analysis techniques that have already been developed and employed by a large research community. The FieldTrip toolbox supports the data formats of most popular electrophysiological data acquisition systems and shares analysis code with other software packages such as SPM and EEGLAB^18^. In contrast to the host of proprietary programs currently available for the analysis of electrophysiological data, the central tenet of FieldTrip is to provide complete transparency in order to promote a deeper understanding of the analysis techniques and enhance the quality of the scientific work that depends on these techniques. Accordingly, all computer code is fully accessible and the well-defined data structures contain full provenance to facilitate sharing between researchers. Our aim is to utilize these open source features to advance the field of human iEEG by promoting interaction within and across methodologically contiguous research areas (e.g., non-invasive electrophysiology such as EEG or MEG).

### Application of the protocol

Our protocol is especially useful for studying human neural and cognitive processes with intracranial EEG. Human iEEG analysis is uniquely complex because it requires dealing with both neuroanatomical and large electrophysiological data sets. The scope of iEEG encompasses a wide range of basic and clinical research, varying from studies of higher-order cognition^19, 20^ to the localization and understanding of the sources and features of epileptogenic activity^21, 22^. The methodological challenges that iEEG researchers face can be grouped into obstacles that are common to most empirical work and obstacles that are study-specific. This protocol aims to resolve the former, while providing adequate support and flexibility for the latter.

### Advantages and limitations of the protocol

The main advantages of our protocol are that it (i) guides the researcher from the multitude of raw intracranial data files to integrated observations, in a fast and efficient way, (ii) is directly integrated with a comprehensive and open source hub for electrophysiological data analysis, (iii) can be readily adapted and automated, (iv) is completely transparent and (v) produces reproducible workflows and data sets that can be easily shared and generalized to other research modalities. The main limitation is that the MATLAB command line interface requires some basic programming knowledge, which may require more initial learning as compared to the execution of computer commands through a (black box) graphical user interface. However, the use of computer commands can be relatively easily mastered by virtue of using this protocol, paving the way for batch scripting in order to efficiently deal with repeated analyses within and across subjects and, ultimately, for a deeper understanding of the underlying algorithms.

Human intracranial data sets are approached from various angles and come in different shapes and sizes, so it is critical for a protocol to strike the right balance between efficiency and flexibility. This need is further amplified by the relatively unique nature of intracranial data, typically imposing greater demands on alternative options and strategies in the analysis than non-invasive data recorded with more standardized hard- and software in dedicated laboratory settings. Besides providing a quick guide to interpretable results, our protocol allows for easy switching between methods to accommodate different cases and situations. By changing a single parameter at execution, one can for instance readily apply a different fusion cost function or filter setting. Utilizing this versatility should not negatively impact continuation with the protocol. In fact, the full and automatic provenance, in combination with the systematic file naming, encourages adapting to the circumstances by alleviating potential concerns regarding oversight and reproducibility.

The spatiotemporal precision of intracranial EEG provides a unique window on neural processing. The size and dimensions of this window, however, may grow disproportionally large with certain types of analyses, complicating the overall interpretability of the data. Starting from the two dimensions of the raw neural signal (channels and time), a time-frequency analysis, for instance, results in 3 dimensions in the output (power as function of channel, time and frequency), whereas between-channel connectivity analysis expands the combinatorial space to 4 dimensions. Our protocol addresses this dimensionality issue and illustrates how the interactive manipulation of anatomically informed graphical representations of the neural data facilitates the inspection of the multi-dimensional outcome of an iEEG analysis, taking maximum advantage of the groundwork laid by the integrated processing of the anatomical and functional data.

### Integration with FieldTrip

In addition to the complete transparency that comes with an open source toolbox, the integration with FieldTrip provides unique benefits to iEEG researchers by allowing them to build on algorithms for reading in raw data of various formats, data preprocessing, event-related potential analysis, spectral analysis, source modeling, connectivity analysis, classification, real-time data processing, and statistical inference. Applied to human iEEG data, these methods permit characterization of neural information flow with a level of detail inaccessible to noninvasive techniques. Additionally, invasive and non-invasive human electrophysiology can be directly overlaid using very similar analysis pipelines for an integrative perspective of neural processing, or a comparison of MEG/EEG source reconstruction methods with iEEG.

The open source development model allows for a relatively easy extension of the protocol. For instance, several techniques exist to compensate for electrode displacement due to the “brain shift” phenomenon explained below^11, 12, 23–30^. Given different strengths and weaknesses, these techniques may need to be evaluated on a case-by-case basis. FieldTrip’s modular architecture facilitates developers to incorporate new techniques and users to subsequently employ those techniques by virtue of changing a single parameter at function call. In a similar vein, the protocol can be extended to a number of exciting new research areas. These include single-and multiunit recordings, ‘NeuroGrid’ recordings^31^, wireless ‘Neural Dust’ recordings^32^, (deep) brain stimulation^33, 34^, and multimodal imaging^12^. Supported by a growing community of developers committed to the ongoing push to improve data analysis methods, we will coordinate with these new electrophysiological endeavors and continue sharing analysis code with other software packages.

### Compatibility with FreeSurfer

The protocol is compatible with the freely available software package FreeSurfer^35^. Although optional, processing of the anatomical MRI with FreeSurfer (Step 6) offers several advantages for subsequent analysis and data interpretation. Processing the MRI with FreeSurfer results in the creation of a cortical mesh, consisting of approximately equally sized triangles that form a topological sphere for each of the cerebral hemispheres. This cortical mesh is particularly convenient for an anatomically realistic representation of the electrophysiological data on the neocortex (e.g., bottom center in Fig. 1). A smoothed version of the extracted cortical surface can be used in the compensation for electrode displacement due to brain shift (Step 22). Moreover, FreeSurfer automatically registers the subject’s brain to a template brain on the basis of its cortical gyrification pattern, an aspect of brain structure that remains difficult to accurately normalize using volume-based registration techniques due to its complexity and variability across subjects^36, 37^. Our protocol uses the resulting surface registration maps to link electrode positions to their template homologues (Step 29). Finally, FreeSurfer-generated atlases are convenient for representations of neural and anatomical data for a single subject (Step 52), since they are defined in native subject space. Other supported atlases are defined in standardized (e.g., MNI) space and require the added step of transforming electrode positions to that space.

### Human intracranial data

Anatomical images, typically MRI and CT scans, are used as part of the epilepsy diagnostic and surgical procedures. A pre-implant MRI shows the anatomy of the head including the brain and is used to identify structural abnormalities. An MRI is also instrumental in guiding SEEG electrode implantation subsequent to the clinical decision to record intracranially. A post-implant CT shows high-intensity objects such as the electrodes and skull but lacks details of brain anatomy. To obtain knowledge of an electrode’s location in relation to the brain’s anatomy, the two scans have to be fused.

Following fusion of the pre- and postoperative anatomical images, electrodes that have been surgically placed on the cortical surface occasionally appear “buried” within the cortical tissue, sometimes more than a centimeter deep^38–43^. This electrode displacement is typically due to “brain shift”, the inward sinking of the brain post-implant most commonly observed with electrocorticographic surface grid electrodes. The brain shift reflects tissue displacement, caused by the electrodes themselves, and by subdural fluid loss or accumulation. As noted, the displacement is most pronounced directly below a craniotomy and is usually minimal for implants solely involving burr holes^43^. It is important to account for this brain shift in order to accurately align electrode specific signals with the local cortical anatomy. Several labs have developed realignment techniques to compensate for electrode displacement due to brain shift, reducing localization error to under 3 mm when compared to intraoperative photographs^11, 23–30^. Our protocol currently supports two of these techniques to project electrode grids back to the cortical surface while accounting for a grid’s shape and orientation^23, 30^.

Electrode localization can also be done using post-implant MRIs, although these are not commonly acquired in a clinical setting. These scans show the brain anatomy after electrode implantation, so brain shift is not an issue. In a T1-weighted MRI, electrodes appear dark, due to the magnetic susceptibility artifact. This is generally not an issue for recordings with depth electrodes (SEEG), where the electrodes are visible as dark voids in the higher intensity brain tissue. Electrode grids and strips (ECoG), on the other hand, are placed directly on the cortical surface. This complicates their identification, as the electrodes are surrounded by cerebral spinal fluid, which also appears dark on a T1 scan (but see 2s,^44–46^ for workarounds). Depending on the availability of a post-implant MRI of sufficient quality that clearly shows the electrodes, the CT preprocessing and fusion Steps 9-15 may be left out, and electrode localization may be done on the post-implant MRI. However, if the post-implant MRI is of unsatisfactory quality regarding brain anatomy, for instance due to electrode induced MR signal distortion, we recommend fusing the post-implant MRI with the pre-implant MRI, as if it were a post-implant CT.

Neural recordings are typically part of the ongoing clinical monitoring and come in various file formats. Each data channel represents, as a function of time, the electric potential difference, obtained with either a bipolar or referential electrode scheme. That is, the electrodes are pairwise linked or referenced to a single, common electrode during acquisition. The latter montage has the benefit that the recordings can be easily re-montaged to a more preferred scheme in the offline analysis^47^. The markers or triggers for stimulus onset times and responses are typically recorded simultaneously in a dedicated channel, allowing for precise synchronization of experimental scenarios with the neural recording.

### Overview of the procedure

The protocol is grounded in two parallel but interrelated workflows, as shown in Figure 1. The first workflow entails the processing of anatomical data. Its main activities constitute the preprocessing and fusion of the anatomical images, and electrode placement (Steps 1-19). Secondary activities that are also discussed include cortical surface extraction with FreeSurfer, brain shift compensation, spatial normalization, and anatomical labeling (Steps 6 and 20-33). Generally, the anatomical workflow aims to obtain estimates of the electrode locations in relation to the individual and atlas-based brain anatomy, which is a one-time procedure for each subject. The second workflow focuses on improving the signal-to-noise ratio and extracting the relevant features from the electrophysiological data, while preparing for subsequent analyses. It minimally encompasses the preprocessing of the neural recordings, but may also include follow-up activities such as time-frequency and single-subject or group-level statistical analysis (Steps 34-45). Generally, the specifics of the functional workflow depend ultimately on the clinical or research question at hand and contingencies in the experimental paradigm.

**Figure 1.**
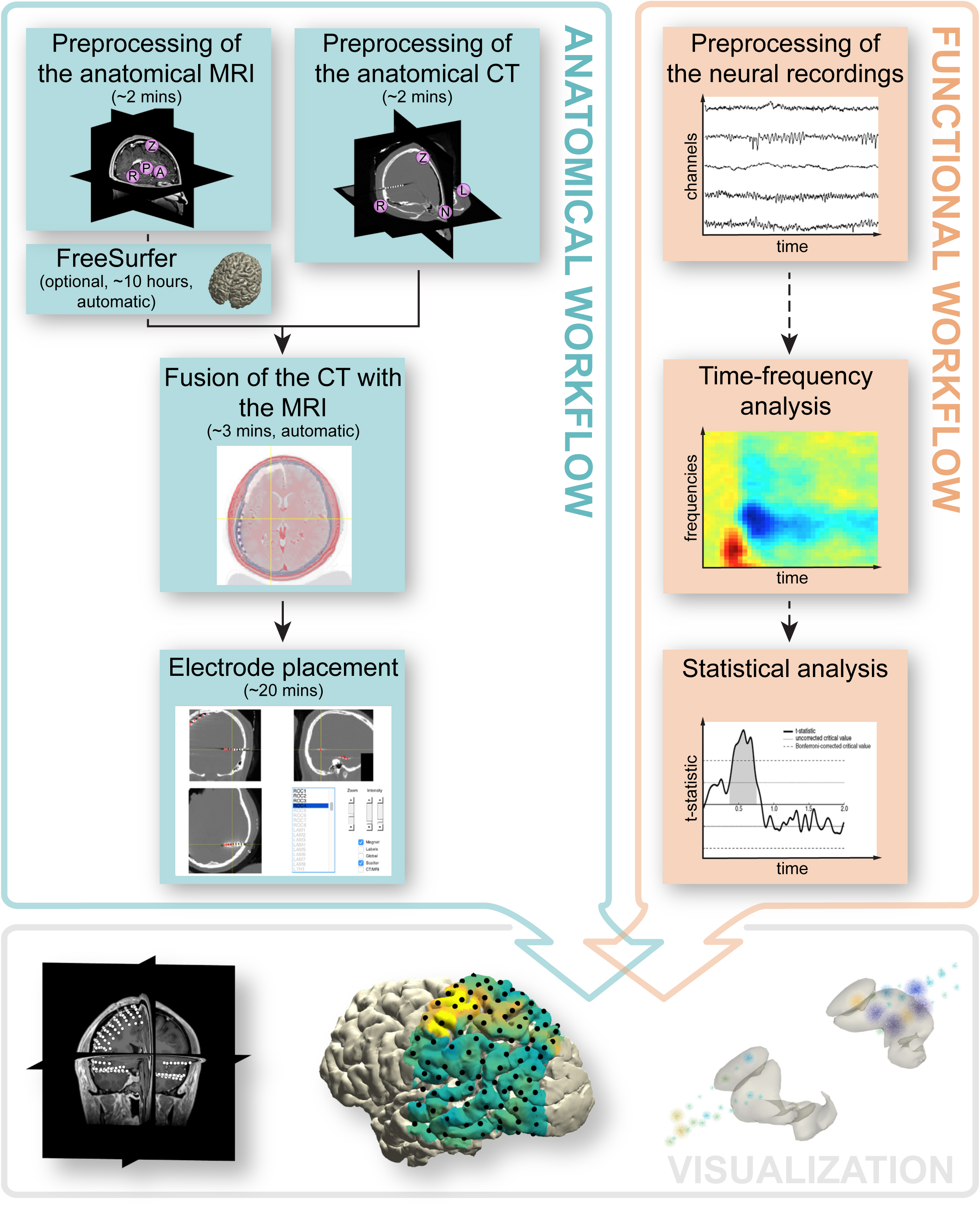
Overview of the procedure. The protocol is grounded in two parallel but interrelated workflows. The anatomical workflow minimally consists of the preprocessing and fusion of the anatomical images and electrode placement. The functional workflow encompasses the preprocessing of the neural recordings, but may also include follow-up activities such as event-related averaging, time-frequency and statistical analysis. The electrode placement activity offers the opportunity to directly link anatomical locations to electrode labels corresponding to the neural recordings, allowing for an early seamless integration of the two workflows to facilitate anatomically informed data exploration and visualization.

The two workflows become intrinsically connected for the first time during the electrode placement activity (Step 17), which offers the opportunity to directly link anatomical locations to electrode labels corresponding to the neural recordings. This activity involves an interactive electrode placement tool designed for efficient yet precise identification of electrodes in even the most challenging cases. The integration of the two workflows culminates in an interactive and anatomically informed data exploration tool and the ability to represent functional and epileptiform neural activity overlaid on cortical and subcortical surface models, in figure or video format (Steps 46-56).

### Implementation and adaptation of the procedure

All implementations run on a single universal platform (MATLAB, except for FreeSurfer) to support relatively easily automated procedures for dealing with repeated analyses within and across subjects. We recommend that the user construct a single script for a single subject by incrementally copy-pasting code from this protocol into the MATLAB editor (Supplementary File 1), and evaluating segments of that script in the MATLAB command window. Once the script produces satisfactory results, it can be converted into a batch analysis by breaking it into separate components. By looping around the separate components for all subjects, the entire analysis pipeline for all subjects in a study can easily be executed and intermediate results can be saved and evaluated.

The whole batch can be documented and shared, or re-evaluated with different parameter settings as appropriate. By virtue of changing single parameters at a function call, one can for instance readily alternate between various fusion, localization, projection, normalization, filtering, re-montaging, and spectral estimation algorithms to accommodate different cases and situations. The output data structures are kept consistent across the different algorithms, and the parameters to the used algorithm are appended to allow for access to the full data provenance at any level of the analysis pipeline (Box 1).

### Experimental design

The example iEEG data set was acquired at the Medical Center of the University of California, Irvine. The Office for the Protection of Human Subjects of the University of California, Berkeley, approved the study and the subject gave informed consent. The data set includes a pre-implant MRI, a post-implant CT, a post-implant MRI, and neural recordings from 96 ECoG and 56 SEEG electrodes that were implanted as part of the preparation for the epilepsy surgery (see Materials). The neural data were recorded in the context of an experiment that required the patient to press a button with the right hand when hearing a target tone. The original data set (after defacing the imaging data with *ft_defacevolume)* and the processed results are available for download from ftp://ftp.fieldtriptoolbox.org/pub/fieldtrip/tutorial/SubjectUCI29.zip. Raw DICOM images and recording files are not shared to protect the subject’s identity.

We choose this iEEG data set for three reasons. First, it contains neural recordings from both cortical grid (ECoG) and stereotactically inserted depth electrodes (SEEG), requiring strategies for dealing with each type as well as their combination in the analysis. Second, the pre-implant MRI is not of the best quality (a contrast agent was used), electrodes of adjacent cortical grids have seemingly merged with one another in the post-implant CT, and there is significant electrode displacement due to a subdural hygroma contributing to brain shift. These issues reflect real world challenges in intracranial data analysis, allowing us to demonstrate the application of our protocol to non-ideal data. Finally, the experimental paradigm is simple enough to need no further explanation, yet requires performing all the fundamental steps underlying the analysis of intracranial data recorded using a more complex experimental paradigm^19, 48^. We demonstrate the analysis of task-related high-frequency-band activity (~70 to 150 Hz), a prominent neural signature in intracranial data that has been associated with neuron population level firing rate^5, 49–52^. Many other supported analyses such as event-related potential analysis, connectivity analysis, and statistical analysis have been described in detail elsewhere^53–55^.

## MATERIALS

### Anatomical images

- Pre-implant T1-weighted MRI (Magnetic Resonance Image, Siemens 3T TrioTim).
- Post-implant CT (Computerized Tomography, Philips iCT 256).
- Post-implant T1-weighted MRI (Magnetic Resonance Image, Siemens 1.5T Avanto). This scan is not used in the procedure but nevertheless included for completeness.

### Neural recordings

- 64-contact cortical grid with left parietal coverage (Integra, 8 x 8 layout, 10 mm inter-electrode spacing, labels have a LPG prefix)
- 32-contact cortical grid with left temporal coverage (Integra, 4 x 8 layout, 10 mm inter-electrode spacing, labels have a LTG prefix)
- 8-contact linear depth electrode targeting left amygdala (Ad-Tech, 5 mm inter-electrode spacing, labels have a LAM prefix)
- 8-contact linear depth electrode targeting left hippocampus head (Ad-Tech, 5 mm inter-electrode spacing, labels have a LHH prefix)
- 8-contact linear depth electrode targeting left hippocampus tail (Ad-Tech, 5 mm inter-electrode spacing, labels have a LTH prefix)
- 8-contact linear depth electrode targeting right amygdala (Ad-Tech, 5 mm inter-electrode spacing, labels have a RAM prefix)
- 8-contact linear depth electrode targeting right hippocampus head (Ad-Tech, 5 mm inter-electrode spacing, labels have a RHH prefix)
- 8-contact linear depth electrode targeting right hippocampus tail (Ad-Tech, 5 mm inter-electrode spacing, labels have a RTH prefix)
- 8-contact linear depth electrode targeting right occipital cortex (Ad-Tech, 5 mm inter-electrode spacing, labels have a ROC prefix)
- All neural recordings were acquired using a Nihon Kohden recording system with a JE-120A amplifier (Nihon Kohden Corporation, Tokyo, Japan), analog-filtered above 0.01 Hz, and digitally sampled at 5 KHz

### Software

- MATLAB environment (MathWorks, Natick, MA; installation and licensing through http://www.mathworks.com)
- FieldTrip toolbox (Box 1, freely available at http://www.fieldtriptoolbox.org)
- FreeSurfer software suite for cortical surface extraction (optional; freely available at http://www.freesurfer.net)

### Supported anatomical data formats

- AFNI (*.head, *.brik)
- Analyze (*.img, *.hdr)
- ANT (*.mri)
- DICOM (*.dcm, *.ima)
- FreeSurfer (*.mgz, *.mgh)
- MINC (*.mnc)
- NIfTI (*.nii, *.nii.gz)

### Supported electrophysiological data formats

- Anywave (*.ah5)
- BCI2000 (*.dat)
- BESA (*.besa)
- Blackrock (*.nev, *.ns#)
- Cambridge Electronic Design (*.smr)
- European Data Format (*.edf)
- GTec (*.mat, *.hdf5)
- Micromed (*.trc)
- Neuralynx (*.ncs, *.nse, *.nts, *.nst, *.ntt, *.nev)
- Neuromag (*.fif)
- Neuroscope (*.eeg, *.dat, *.xml)
- Nihon Kohden (*.m00)
- Plexon (*.ddt, *.nex, *.plx)
- and various EEG, MEG, NIRS, and eye-tracker data formats

## PROCEDURE

1| Specify the subject ID. This ID will be used in the file naming, in addition to information about the type of data (e.g., MRI, CT), the coordinate system it is in (e.g., ACPC, MNI), and the process(es) that were applied to it (e.g., f for fusion). For example, a CT scan that is aligned to the ACPC coordinate system and that has just been fused with the anatomical MRI is written out to file as subjID_CT_acpc_f.nii.

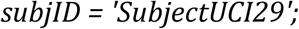

### Preprocessing of the anatomical MRI, TIMING ~2 min

2| Import the anatomical MRI into the MATLAB workspace using *ft_read_mri.* The MRI comes in the format of a single file with an .img or .nii extension, or a folder containing a series of files with a .dcm or .ima extension (DICOM; Supplementary File 2 may aid in the search and visualization of a DICOM series).

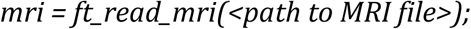

3| Determine the native orientation of the anatomical MRI’s left-right axis using *ft_determine_coordsys* (Box 2 and Supplementary Video 1).

#### CRITICAL STEP

To correctly fuse the MRI and CT scans at a later step, accuracy in demarcating the right hemisphere landmark in the following step is important for avoiding an otherwise hard to detect flip of the scan’s left and right orientation.

4| Align the anatomical MRI to the ACPC coordinate system^56^, a preferred convention for the FreeSurfer operation optionally used in a later step. In this coordinate system, the origin (coordinate [0,0,0]) is at the anterior commissure (AC), the Y-axis runs along the line between the anterior and posterior commissure (PC), and the Z-axis lies in the midline dividing the two cerebral hemispheres. Specify the anterior and posterior commissure, an interhemispheric location along the midline at the top of the brain, and a location in the brain’s right hemisphere. If the scan was found to have a left-to-right orientation in the previous step, the right hemisphere is identified as the hemisphere having larger values along the left-right axis. Vice versa, in a right-to-left system, the right hemisphere has smaller values along that axis than its left counterpart (Supplementary Video 2).

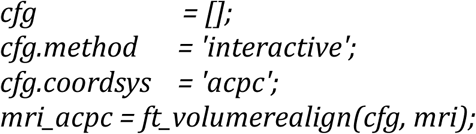

5| Write the preprocessed anatomical MRI out to file.

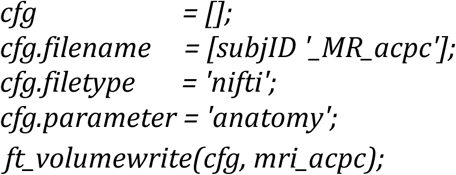

### Cortical surface extraction with FreeSurfer (optional), TIMING ~10 hrs, automatic

6| Execute FreeSurfer’s *recon-all* functionality from the Linux or MacOS terminal (Windows via VirtualBox), or from the MATLAB command window as below. This set of commands will create a folder named ‘freesurfer’ in the subject directory, with subdirectories containing a multitude of FreeSurfer-generated files.

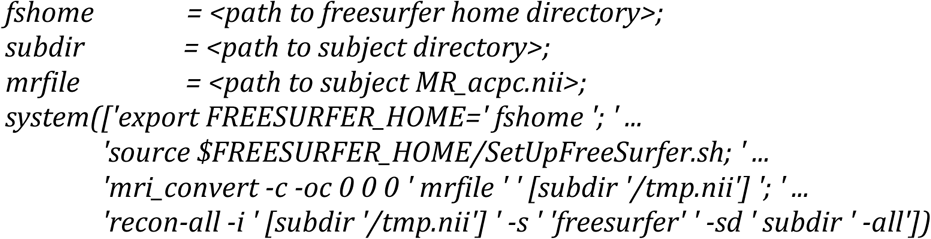

#### PAUSE POINT

FreeSurfer’s fully automated segmentation and cortical extraction of the anatomical MRI currently may take up 10 hours or more. For tutorial purposes, the example data set contains the output from FreeSurfer, a folder named ‘freesurfer’, for continuation with the protocol.

7| Import the extracted cortical surfaces into the MATLAB workspace and examine their quality. Repeat the following code using *rh.pial* to visualize the pial surface of the right hemisphere.

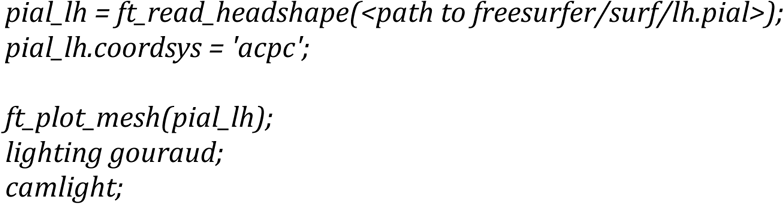

## TROUBLESHOOTING

8| Import the FreeSurfer-processed MRI into the MATLAB workspace for the purpose of fusing with the CT scan at a later step, and specify the coordinate system to which it was aligned in Step 4.

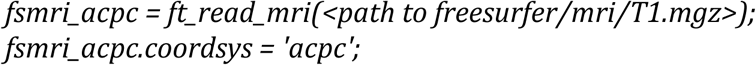

### Preprocessing of the anatomical CT, TIMING ~2 min

9| Import the anatomical CT into the MATLAB workspace. Similar to the MRI, the CT scan comes in the format of a single file with an .img or .nii extension, or a folder containing a series of files with a .dcm or .ima extension (Supplementary File 2 may aid in the search and visualization of a DICOM series).

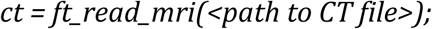

10| In case this cannot be done on the basis of knowledge of the laterality of electrode implantation, determine the native orientation of the anatomical CT’s left-right axis using *ft_determine_coordsys,* similarly to how it was done with the anatomical MRI in Step 3 (Box 2 and Supplementary Video 1).

#### CRITICAL STEP

To correctly fuse the MRI and CT scans at a later step, accuracy in demarcating the right and left preauricular landmark in the following step is important for avoiding an otherwise hard to detect flip of the scan’s left and right orientation.

11| Align the anatomical CT to the head surface coordinate system by specifying the nasion (at the root of the nose), left and right preauricular points (just in front of the ear canals), and an interhemispheric location along the midline at the top of the brain (Supplementary Video 3). The CT scan is initially aligned to the head surface coordinate system, given that the ACPC coordinate system used for the MRI relies on neuroanatomical landmarks that are not visible in the CT.

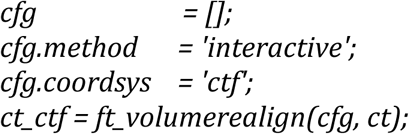

12| Automatically convert the CT’s coordinate system into an approximation of the ACPC coordinate system, the same system the anatomical MRI was aligned to.

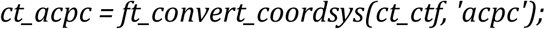

### Fusion of the CT with the MRI, TIMING ~3 min

13| Fuse the CT with the MRI, a necessary step to link the electrode locations in the anatomical CT to their corresponding locations in the anatomical MRI^57^,^58^. Given that both scans are from the same subject and their common denominator is the skull, a rigid body transformation suffices for their alignment under normal circumstances (the default technique when using the SPM-method in FieldTrip).

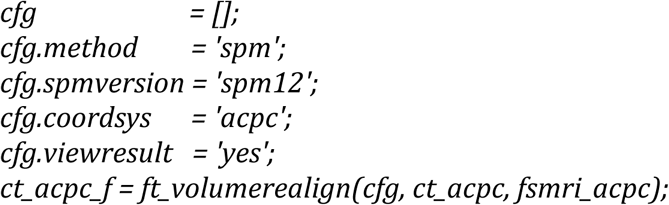

14| Carefully examine the interactive figure that is produced after the coregistration is completed, showing the MRI and fused CT superimposed. A successful fusion will show tight interlocking of CT-positive skull (in blue) and MRI-positive brain and skin tissue (in red).

#### CRITICAL STEP

Accuracy of the fusion operation is important for correctly placing the electrodes in anatomical context in a following step.

## TROUBLESHOOTING

15| Write the MRI-fused anatomical CT out to file.

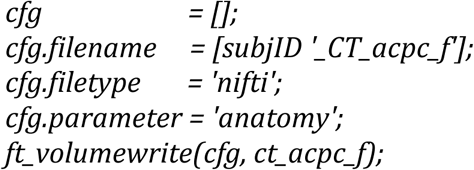

### Electrode placement, TIMING ~15 min

16| Import the header information from the recording file, if possible. By giving the electrode labels originating from the header as input to *ft_electrodeplacement* in the next step, the labels will appear as a to-do list during the interactive electrode placement activity. A second benefit is that the electrode locations can be directly assigned to labels collected from the recording file, obviating the need to sort and rename electrodes to match the electrophysiological data.

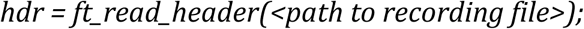

17| Localize the electrodes in the post-implant CT with *ft_electrodeplacement*, shown in Figure 2. Clicking an electrode label in the list will directly assign that label to the current crosshair location (Supplementary Video 4). Several in-app features facilitate efficient yet precise navigation of the anatomical image, such as a zoom mode, a magnet option that transports the crosshair to the nearest weighted maximum with subvoxel accuracy (or minimum in case of a post-implant MRI), and an interactive three-dimensional scatter figure that is linked to the two-dimensional volume representations. Furthermore, passing on the pre-implant MRI, *fsmri_acpc*, to *ft_electrodeplacement* allows toggling between CT and MRI views for the identification of specific electrodes based on their anatomical location. Generally, electrode #1 is the electrode farthest away from the craniotomy or burr hole in case of depths and single-row strips. Careful notes taken during surgery and recording are critical for determining the numbering of grid and multi-row strip electrodes.

**Figure 2.**
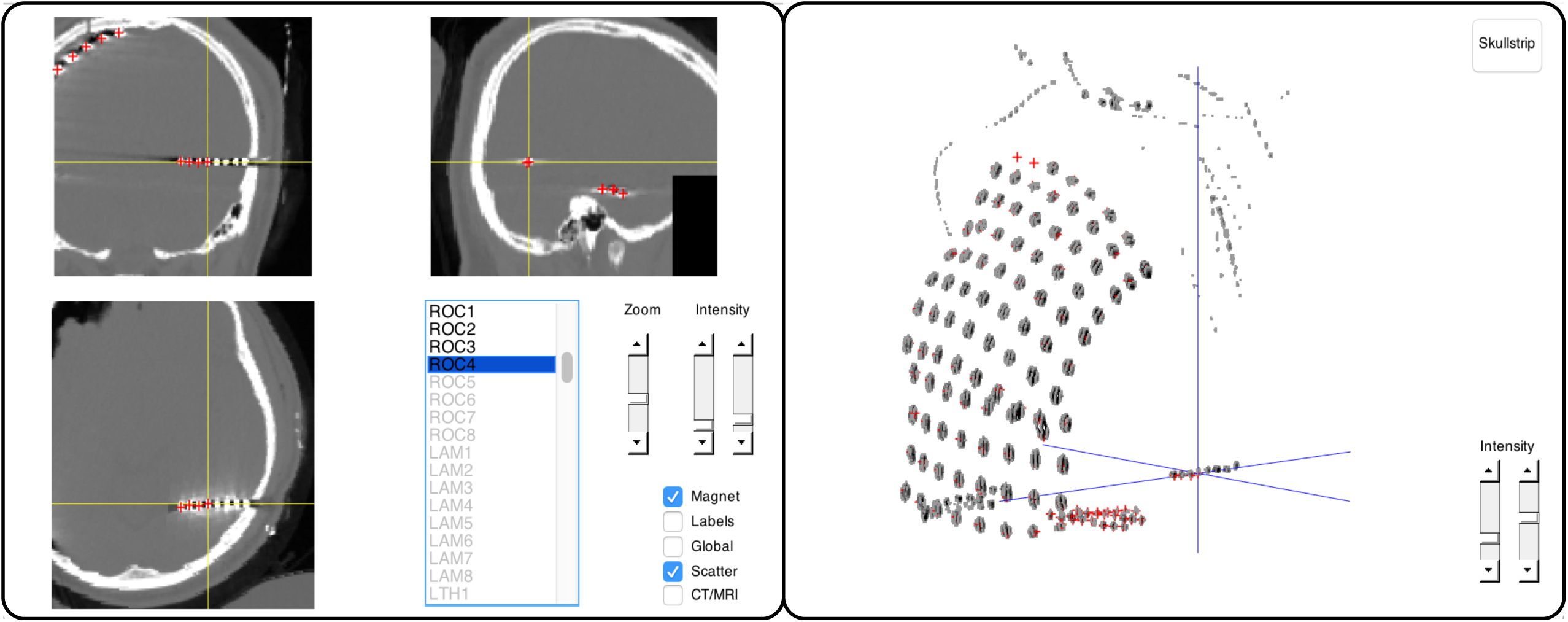
Interactive electrode placement. Clicking an electrode label in the main panel on the left will directly assign that label to the current crosshair position in the CT scan. Several features facilitate precise navigation of the anatomical CT, such as a zoom mode, a magnet option that transports the crosshair to the nearest weighted maximum (or minimum in case of a post-implant MRI), and the interactive three-dimensional scatter figure shown on the right.

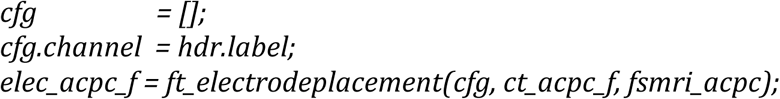

18| Examine whether the variables in resulting electrode structure *elec_acpc_f* match the recording parameters, e.g., the number of channels stored in the *label* field. The electrode and channel positions are stored in the *elecpos* and *chanpos* fields, respectively. The *elecpos* field contains the original electrode positions. With exception of possible brain shift compensation, this field is not adjusted. The channel positions in the *chanpos* field are initially identical to the electrode positions but may be updated to accommodate offline adjustments in channel combinations, i.e. during re-montaging. For bipolar iEEG data, the best considered channel position is in between the two corresponding electrode positions. The chanpos field is used for overlaying the neural data on (sub-)cortical models during data visualization. The *tra* field is a matrix with the weight of each electrode into each channel, which at this stage merely is an identity matrix reflecting one-to-one mappings between electrodes and channels.

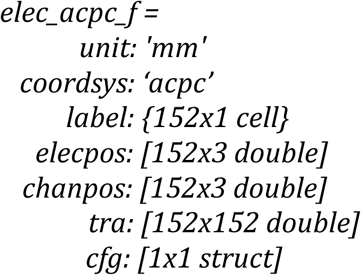

19| Save the resulting electrode information to file.

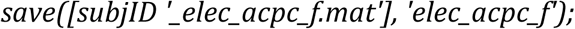

### Brain shift compensation (optional for cortical grids and strips), TIMING ~5 min

21| In case of “brain shift”, a displacement of brain tissue and electrodes postimplant, realignment of electrode grids to the preoperative cortical surface may be necessary. To prevent electrodes from being incorrectly placed in the nearby cortical sulci during back-projection, create a smooth hull around the cortical surface generated by FreeSurfer^59^.

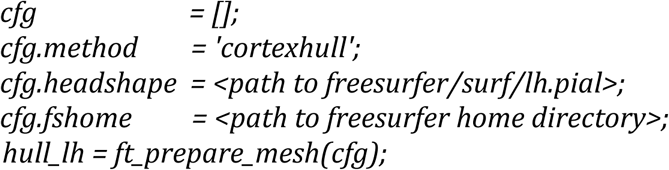

21| Save the hull to file.

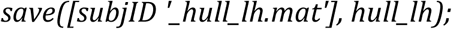

22| Project the electrode grids to the surface hull of the implanted hemisphere. Given that different grids can move independently from one another and that the projection algorithm specified in *cfg.warp* considers the global electrode configuration of a grid^30^, it is recommended to realign electrode grids individually by running separate realignment procedures for each grid. Here, we realign the electrodes of the left parietal grid followed by the electrodes of the left temporal grid (LPG and LTG respectively) and store the updated grid electrode information in a new variable together with the unaltered coordinates of the depth electrodes.

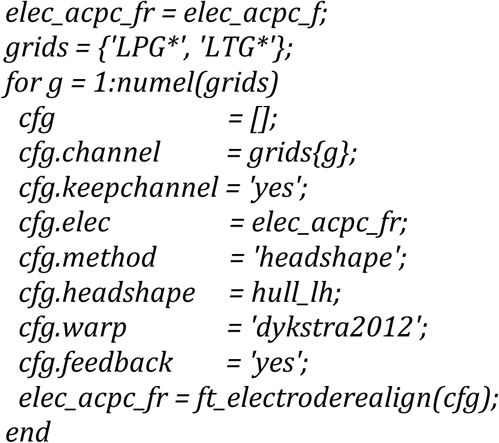

23| Visualize the cortex and electrodes together and examine whether they show expected behavior (Fig. 3).

**Figure 3.**
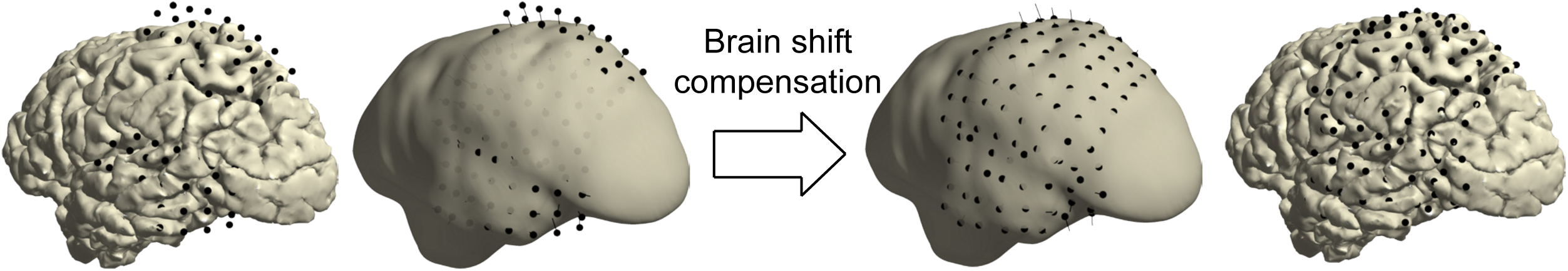
Brain shift compensation. In some patients, compensation for electrode displacement due to brain shift after implantation may be necessary. In this particular case, a subdural hygroma at the top of the brain caused severe electrode displacement in a direction opposite to the more commonly observed inward shift (left). Realigning electrode grids to the cortical surface can compensate for electrode displacement due to brain shift (right). The thin black lines indicate each electrode’s path from its localized origin on the left to its projected location on the right.

#### CRITICAL STEP

Accuracy of the realignment operation is important for correctly placing the electrodes in anatomical context in a following step.

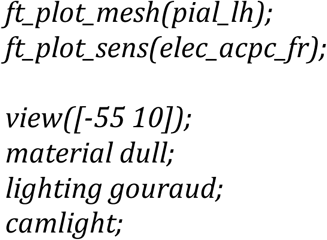

## TROUBLESHOOTING

24| Save the updated electrode information to file.

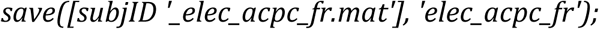

### Volume-based registration (optional), TIMING ~2 min

25| To generalize the electrode coordinates to other brains or MNI-based neuroanatomical atlases in a later step, register the subject’s brain to the standard MNI brain. The volume-based registration technique considers the overall geometry of the brain^60^ and can be used for the spatial normalization of all types of electrodes, whether depth or on the surface.

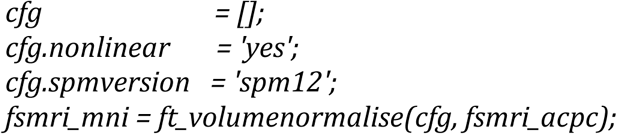

26| Use the resulting deformation parameters to obtain the electrode positions in standard MNI space.

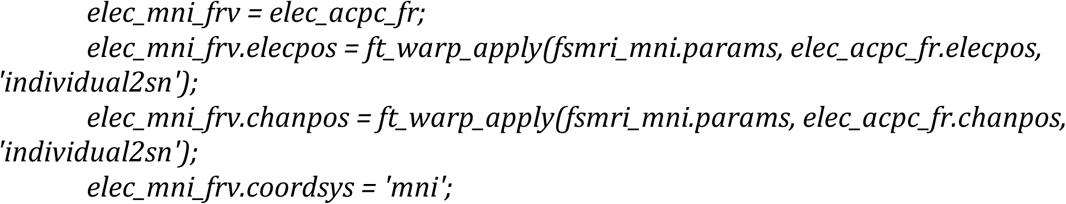

27| Visualize the cortical mesh extracted from the standard MNI brain along with the spatially normalized electrodes and examine whether they show expected behavior (top right in Fig. 4).

**Figure 4.**
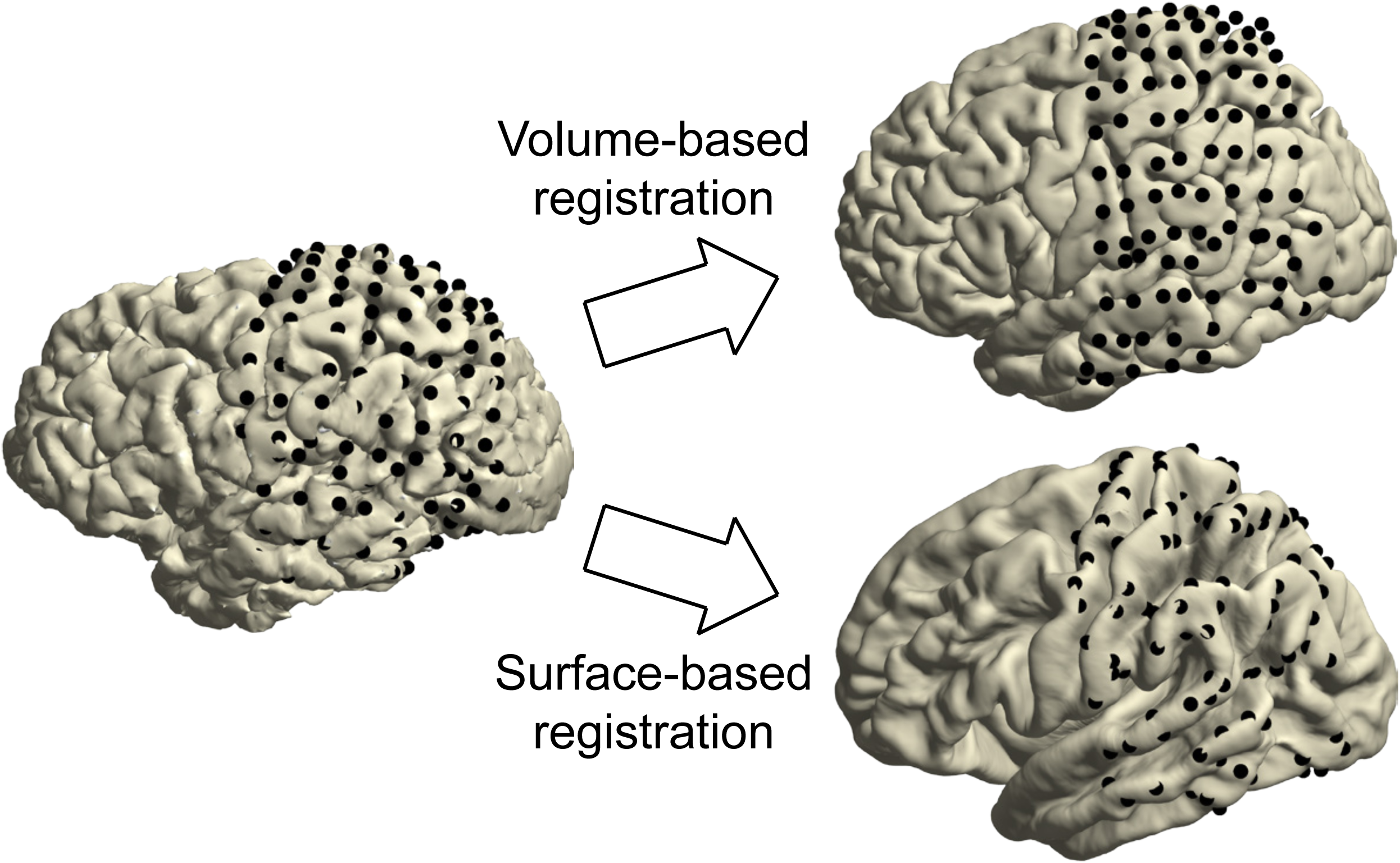
Spatial normalization. On the left are the electrodes on the individual cortical sheet. The top right shows the electrodes on the standard MNI brain after volume-based registration. The bottom right shows the electrodes on FreeSurfer’s fsaverage brain after surface-based registration. Compared to volume-based registration, with surface-based registration the original grid geometry is no longer preserved as electrodes are moved from one brain to another according to the curvature pattern of the cortex.

#### CRITICAL STEP

Accuracy of the spatial normalization step is important for correctly overlaying the electrode positions with a brain atlas in a following step.

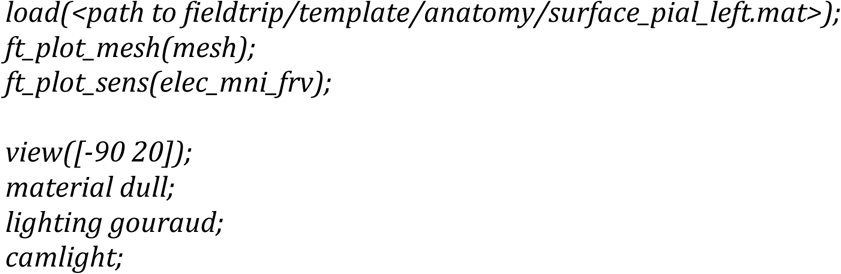

## TROUBLESHOOTING

28| Save the normalized electrode information to file.

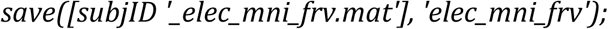

### Surface-based registration (optional for surface electrodes), TIMING ~2 min

29| To generalize the electrode coordinates to other brains in a later step, map the electrodes onto FreeSurfer’s fsaverage brain. The surface-based registration technique solely considers the curvature patterns of the cortex^35^ and thus can be used for the spatial normalization of electrodes located on or near the cortical surface. In the example case, this pertains to all electrodes of the left parietal and temporal grids.

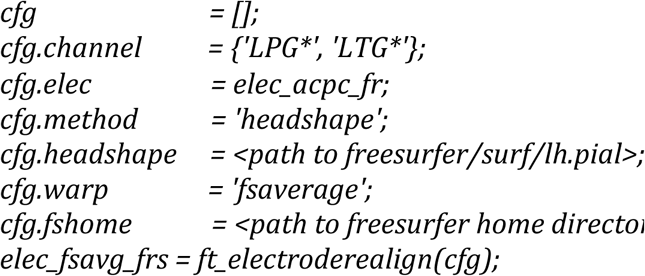

30| Visualize FreeSurfer’s fsaverage brain along with the spatially normalized electrodes and examine whether they show expected behavior (bottom right in Fig. 4).

#### CRITICAL STEP

Accuracy of the spatial normalization step is important for correctly overlaying the electrode positions with a brain atlas in a following step.

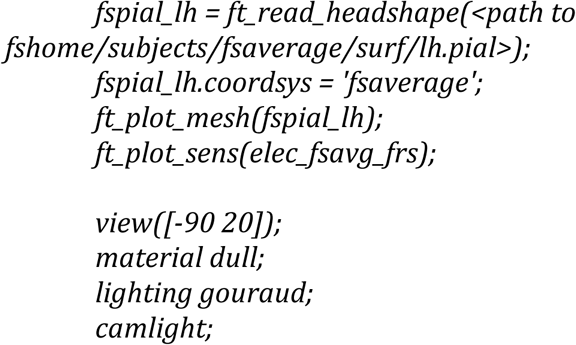

31| Save the normalized electrode information to file.

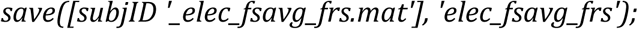

### Anatomical labeling (optional), TIMING ~1 min

32| FieldTrip supports looking up the anatomical or functional labels corresponding to the electrodes in a number of atlases, including the AFNI Talairach Tournoux atlas^61^, the AAL atlas^62^, the BrainWeb data set^63^, the JuBrain cytoarchitectonic atlas^64^, the VTPM atlas^65^, and the Brainnetome atlas^66^, in addition to the subject-tailored Desikan-Killiany and Destrieux atlases produced by FreeSurfer^67, 68^. With exception of the above FreeSurfer-based atlases, these atlases are in MNI coordinate space and require the electrodes to be spatially normalized (Step 25). First, import an atlas of interest, e.g., the AAL atlas, into the MATLAB workspace.

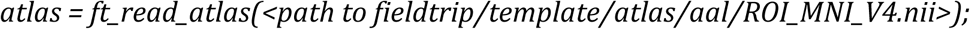

33| Look up the corresponding anatomical label of an electrode of interest, e.g., electrode LHH2, targeting the left hemisphere’s hippocampus. Supplementary File 3 represents a tool that automatically overlays all channels in an electrode structure with all of the above atlases and stores the resulting anatomical labels in an excel table (e.g., SubjectUCI29_electable.xlsx in the zip file).

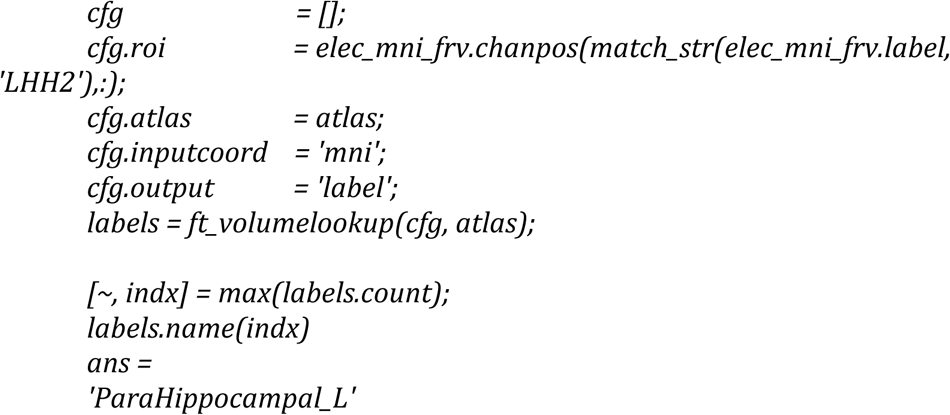

## TROUBLESHOOTING

### Preprocessing of the neural recordings, TIMING ~10 min

34| Define the trials, that is, the segments of data that will be used for further processing and analysis. This step produces a matrix *cfg.trl* containing for each segment the begin and end sample in the recording file. In the case of the example provided in the shared data, the segments of interest begin 400 ms before tone onset, are marked with a ‘4’ in the trigger channel, and end 900 ms thereafter.

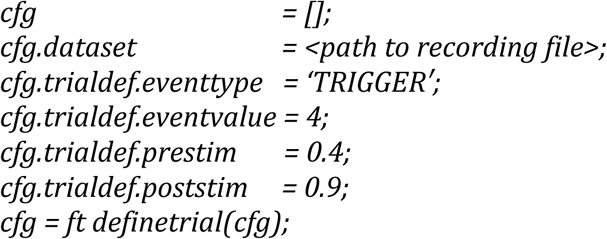

35| Import the data segments of interest into the MATLAB workspace and filter the data for high-frequency and power line noise (see the documentation of *ft_preprocessing* for filtering options).

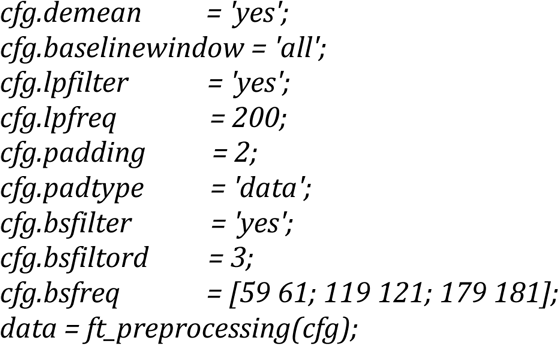

36| Examine whether the variables in the output data structure match the recording and preprocessing parameters, i.e. the sampling rate *fsample*), number of recording channels (*label*), and segmentation into the experiment’s twenty-six trials (*trial,* and their respective time axes in *time*).

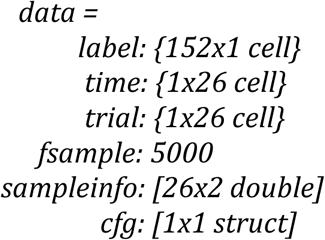

37| Add the *elec* structure originating from the anatomical workflow and save the preprocessed electrophysiological data to file. The advantage of adding the electrode information at this stage is that it will be kept consistent with the neural data going forward, as when applying the same montage used for the neural recordings to the channel positions.

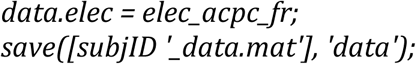

38| Inspect the neural recordings using *ft_databrowser* and identify channels or segments of non-interest, for instance segments containing signal artifacts or (in this case) epileptiform activity. Mark the bad segments by drawing a box around the corrupted signal. Write down the labels of bad channels.

#### CRITICAL STEP

Identifying bad channels is important for avoiding the contamination of other channels during re-montaging in Step 40.

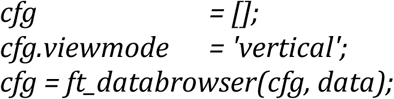

39| Remove any bad segments marked in the above step.

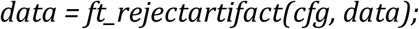

40| Re-montage the cortical grids to a common average reference in order to remove noise that is shared across all channels. Box 3 provides a background on re-montaging. Bad channels noted in Step 38 can be excluded from this step by adding those channels to *cfg.channel* with a minus prefix. That is, *cfg.channel = {‘LPG*’, ‘LTG*’, ‘-LPG1’ }* if one were to exclude the LPG1 channel from the list of LPG and LTG channels.

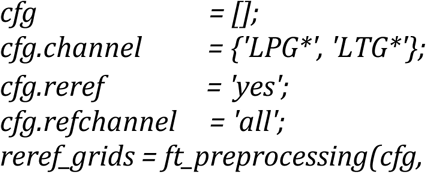

41| Apply a bipolar montage to the depth electrodes. This can be done in a similar manner as in Step 40, but by selecting single channel labels for *cfg.channel* and *cfg.refchannel*. Alternatively, create a more elaborate scheme with *cfg.montage* (see the documentation of *ft_apply_montage*). Here, we combine for each depth electrode shaft the 8 unipolar channels into 7 bipolar channels, using the weights defined in the 7x8 *montage.tra* field. We also create new labels indicating the bipolar origin of the data, e.g., “RAM1-RAM2”, “RAM2-RAM3”, and so on. Note that because we added the *elec* structure to the data in Step 37, the same montage is automatically applied to the channel positions as well, with the resulting *chanpos* field containing the mean locations of all electrode pairs that comprise a bipolar channel.

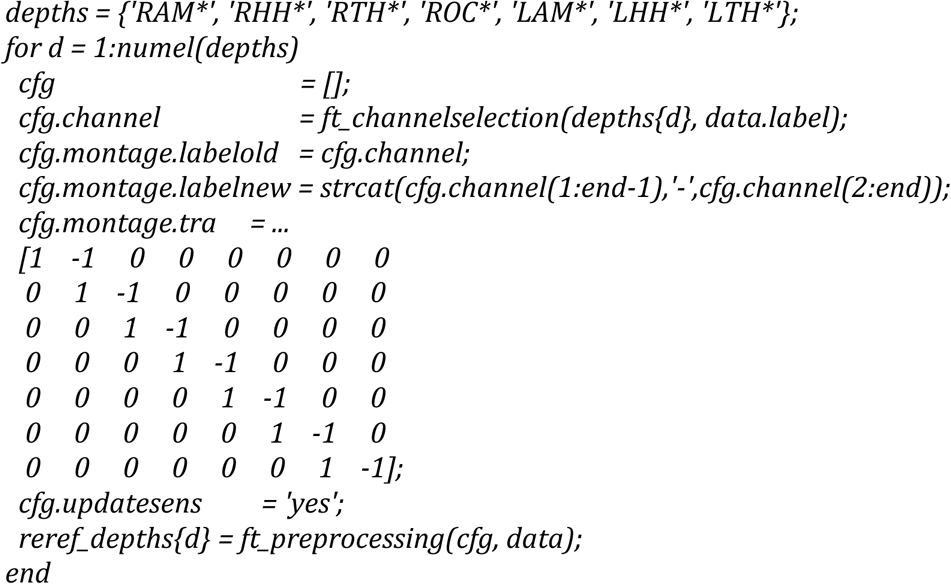

42| Combine the data from both electrode types into one data structure for the ease of further processing.

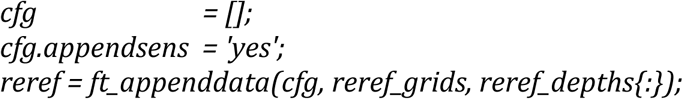

43| Save the re-referenced data to file.

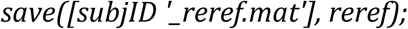

### Time-frequency analysis (optional), TIMING ~2 min

44| Decompose the signal in time and frequency bins. The configuration options *cfg.foi* and *cfg.toi* determine the frequencies and time-points of interest, in this case from 5 to 200 Hz in steps of 5 Hz, and 300 ms prior to tone onset until 800 ms thereafter in steps of 10 ms.

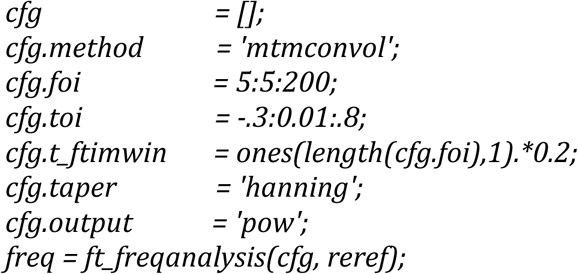

45| Save the time-frequency data to file.

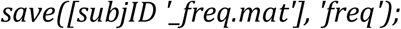

### Interactive plotting, TIMING ~3 min

46| For an anatomically informed exploration of the multidimensional outcome of an analysis, create a layout based on the three-dimensional electrode locations. This layout is a symbolic representation in which the channels are projected on the twodimensional medium offered by paper or a computer screen. The layout is complemented by an automatic outline of the cortical sheet that is specified in *cfg.headshape*. The *cfg.boxchannel* option allows selecting channels whose twodimensional distances are used to determine the plotting box sizes in the following step.

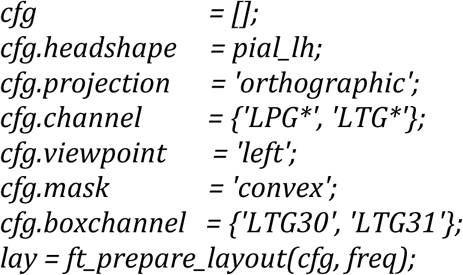

47| Express the time-frequency representation of neural activity at each channel in terms of the relative change in activity from a baseline interval.

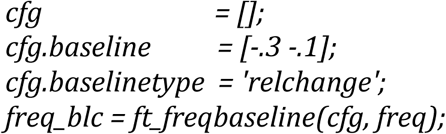

48| Visualize the time-frequency representations overlaid on the two-dimensional layout. The generated figure is interactive, so that selecting a group of channels will launch another figure representing the average time-frequency representation over those channels (Fig. 5). Selecting a certain frequency and time range in that time-frequency representation will launch yet another figure showing the topographical distribution of activity in the selected interval, and so on (Supplementary Video 5).

**Figure 5.**
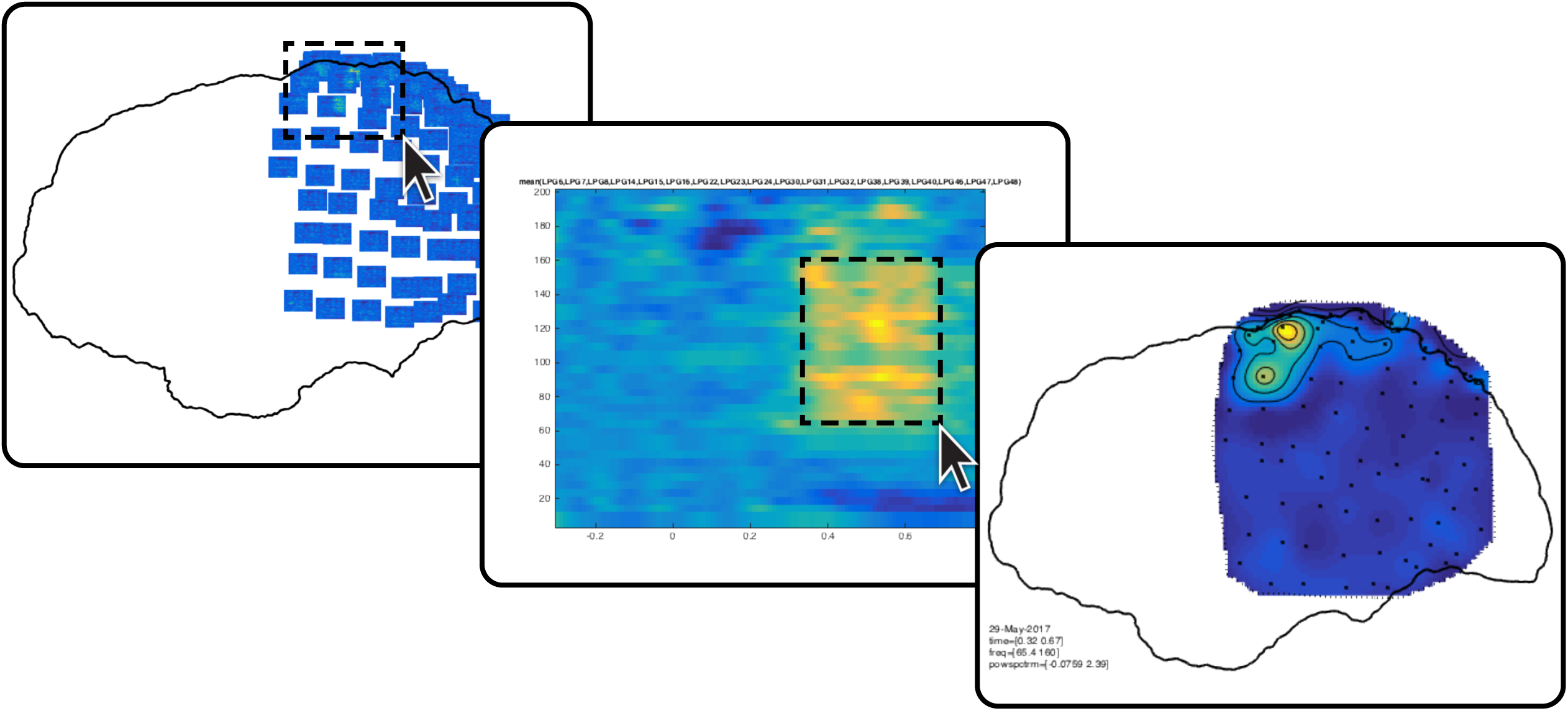
Interactive plotting. Fast browsing through various anatomically informed representations of the neural data can help address the multidimensionality of intracranial EEG data.

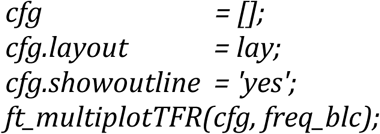

### ECoG data representation, TIMING ~1 min

49| For an anatomically realistic representation of cortical activity, overlay a surface model of the neocortex with the spatial distribution of the high frequency-band activity. First, extract high-frequency-band activity during a time interval of interest.

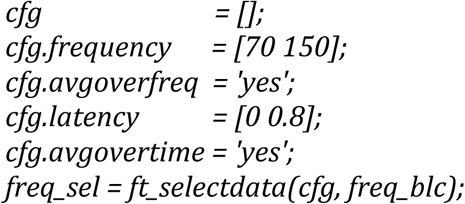

50| Visualize the spatial distribution of high-frequency-band activity on a cortical mesh of the subject’s brain.

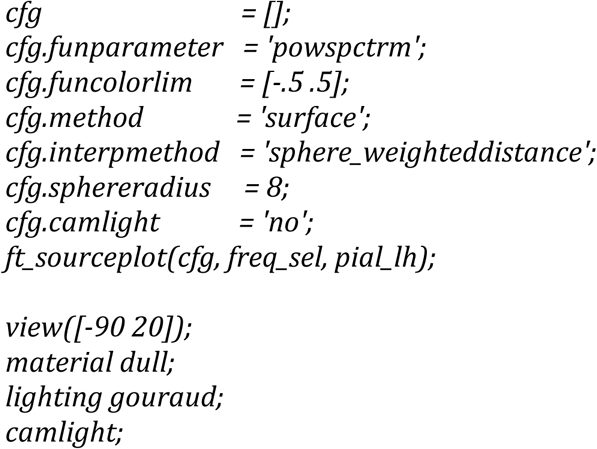

51| Add the electrodes to the figure (Fig. 6). By looping around Steps 49 to 51 while breaking down the time interval of interest specified with *cfg.latency* in consecutive steps, it becomes feasible to observe the spatiotemporal dynamics of neural activity occurring in relation to known experimental structure and behavior (Supplementary Video 6). See *help getframe* for capturing and assembling timelapse movies.

**Figure 6.**
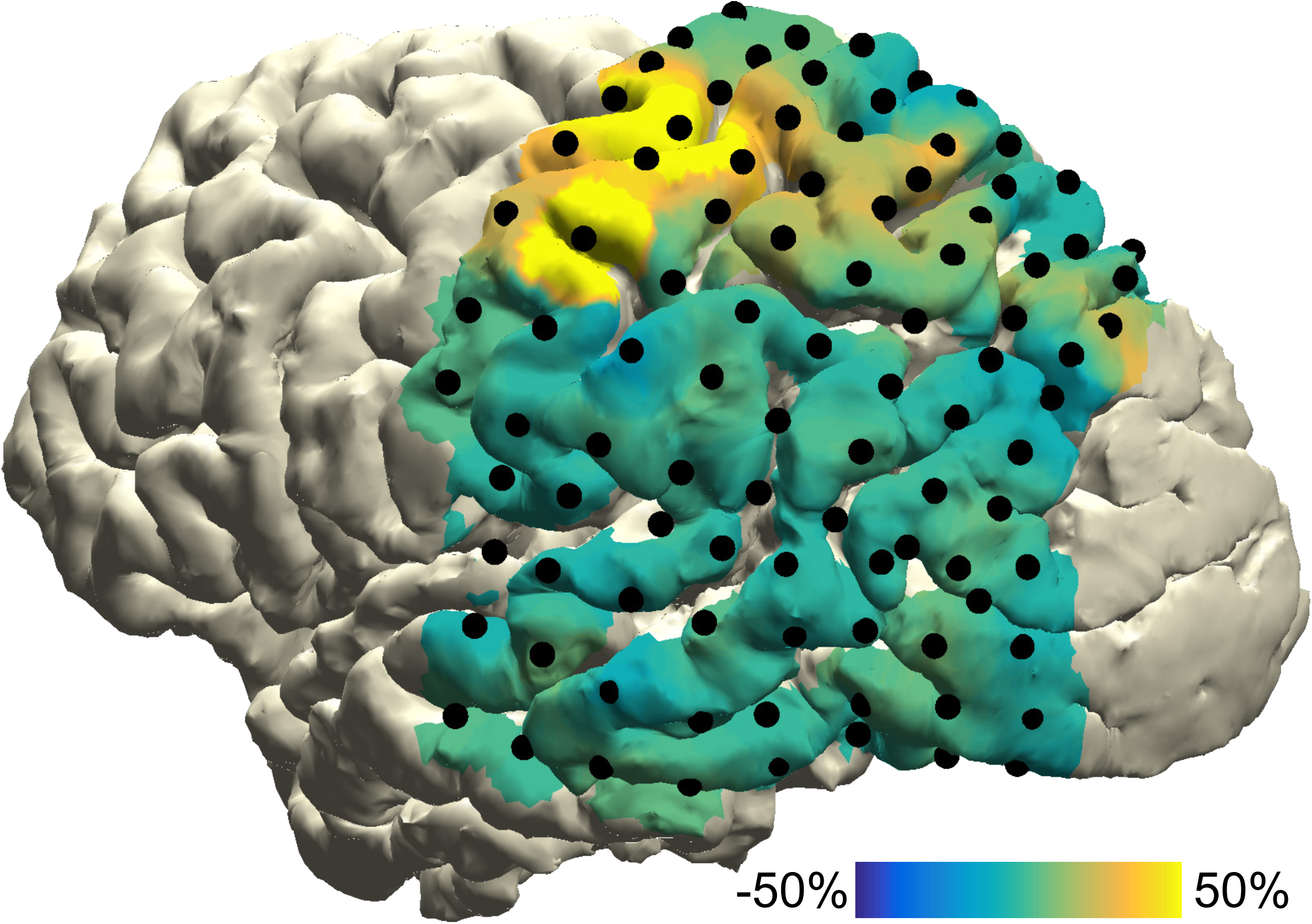
ECoG data representation. Task-induced high-frequency-band activity relative to a baseline interval, plotted on a cortical surface mesh of the subject’s brain.

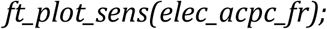

### SEEG data representation, TIMING ~2 min

52| For depth recordings, create an integrated representation of neural activity and anatomy by interpolating neural data from each bipolar channel in a spherical cloud, which can then be overlaid on a surface mesh of any deep brain structure. First, create a volumetric mask of the regions of interest (ROI). Here, we generate a mask for the right hippocampus and amygdala from the cortical parcellation and subcortical segmentation produced by FreeSurfer.

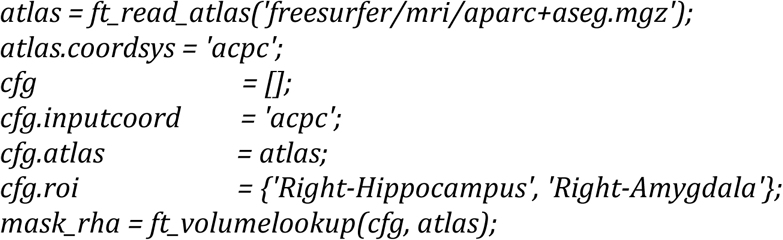

53| Create a triangulated and smoothed surface mesh on the basis of the volumetric masks.

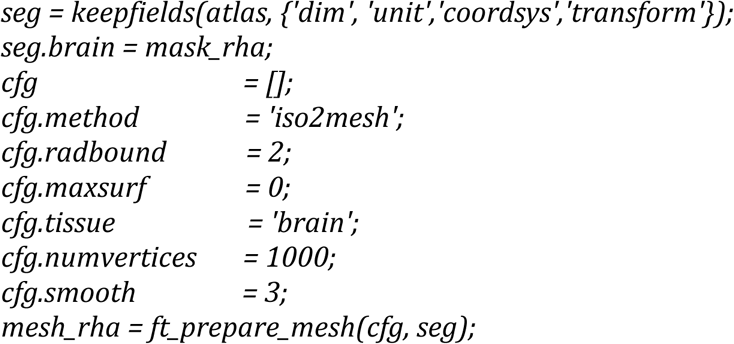

54| Identify the subcortical electrodes of interest.

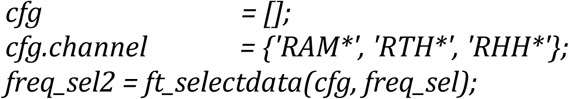

55| Interpolate the high-frequency-band activity in the bipolar channels on a spherical cloud around the channel positions, while overlaying the neural activity with the above mesh. By repeating the current step for neural data corresponding to consecutive time intervals, similarly to the process outlined in Step 51, it becomes feasible to create time-lapse movies of the spatiotemporal dynamics of deep-brain activity (Supplementary Video 7 shows the spatiotemporal evolution of epileptiform activity in a separate subject).

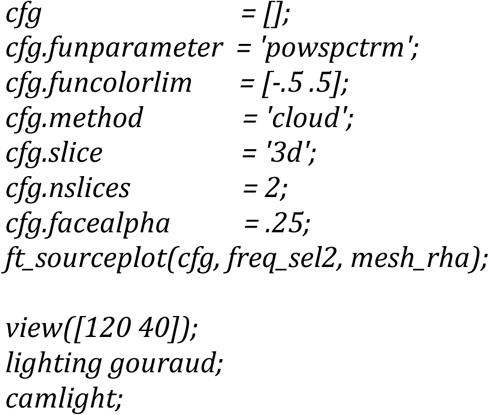

56| To create a more definitive image of the neural activity at particular positions, generate two-dimensional slices through the three-dimensional representations. This combination provides the most complete and integrated representation of neural and anatomical data (Fig. 7).

**Figure 7.**
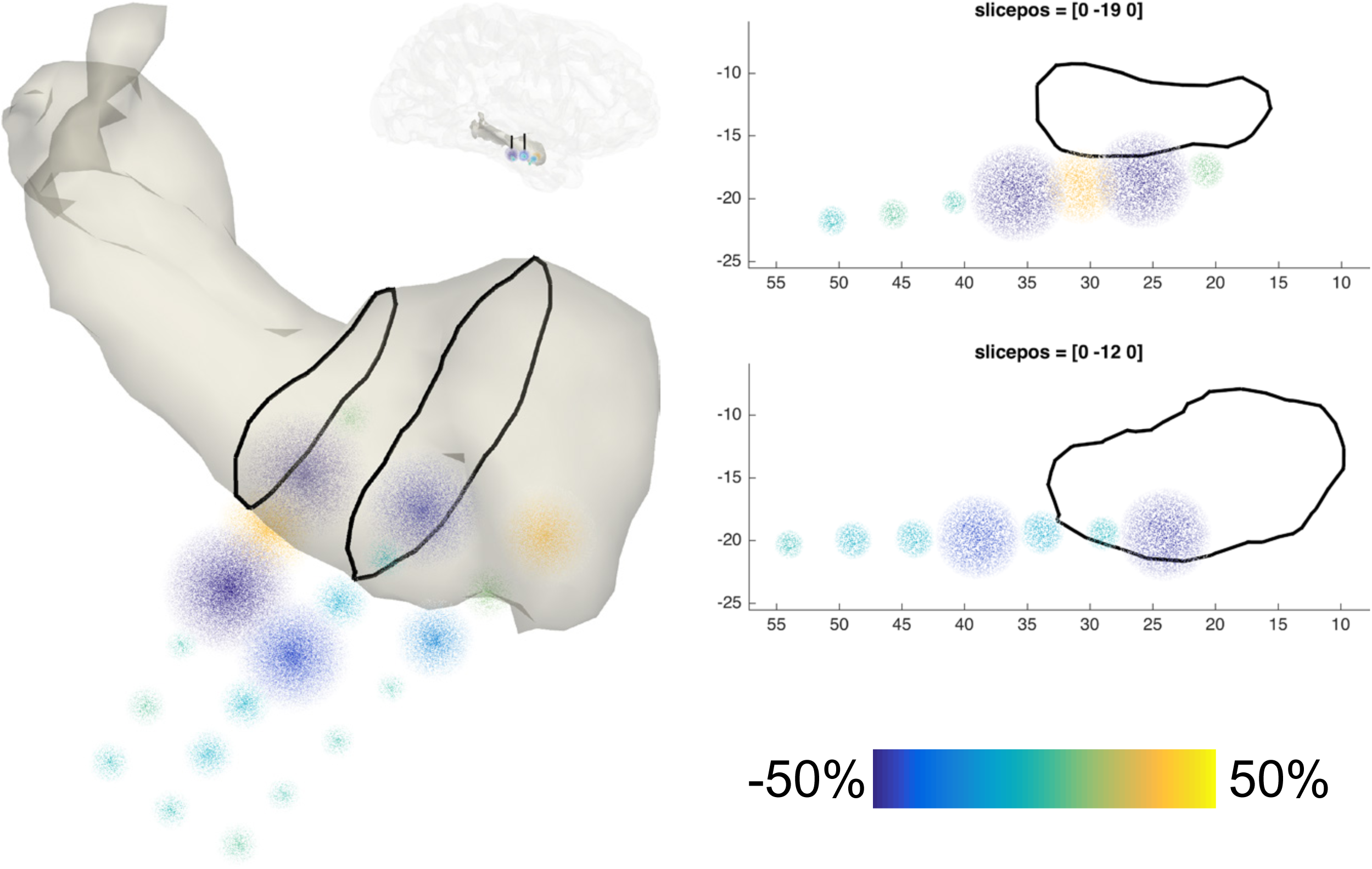
SEEG data representation. Task-induced high-frequency-band activity relative to a baseline interval, plotted as point clouds around a triangulated mesh of the subject’s amygdala and hippocampus in the right hemisphere. The two-dimensional planes on the right correspond to the slices in the image on the left.

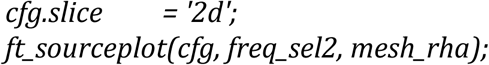

## TIMING

Steps 2-5, Preprocessing of the anatomical MRI: ~2 min
Steps 6-8, Cortical surface extraction with FreeSurfer (optional): ~10 hrs
Steps 9-12, Preprocessing of the anatomical CT: ~2 min
Steps 13-15, Fusion of the CT with the MRI: ~3 min
Steps 16-19, Electrode placement: ~15 min
Steps 21-24, Brain shift compensation (optional): ~5 min
Steps 25-28, Volume-based registration (optional): ~2 min
Steps 29-31, Surface-based registration (optional): ~2 min
Steps 32-33, Anatomical labeling (optional): ~1 min
Steps 34-43, Preprocessing of the neural recordings: ~10 min
Steps 44-45, Time-frequency analysis (optional): ~2 min
Steps 46-48, Interactive plotting: ~3 min
Steps 49-51, ECoG data representation: ~1 min
Steps 52-56, SEEG data representation: ~2 min
Box 2, Coordinate system determination: ~1 min

## ANTICIPATED RESULTS

Upon completion of the protocol, one should obtain an integrated representation of neural and anatomical data. The exact results depend ultimately on the clinical or research question at hand, contingencies in the experimental paradigm, and decisions made during the execution of the protocol. We demonstrated the analysis of spatiotemporal neural dynamics occurring in relation to known experimental structure and relatively simple behavior, namely the pressing of a button with the right hand when hearing a target tone (Fig. 5-7, Supplementary Video 6). However, with small adaptations of the protocol it is feasible to track the spatiotemporal evolution of epileptiform activity with high precision (Supplementary Video 7), or to perform group-level investigations of fine-grained decision-related neural dynamics in human orbitofrontal cortex^48^. A precise fusion of the anatomical images with the electrophysiological data is key to reproducible analyses and findings. Hence, it is important to examine the outcome of any critical step, as we have done in this protocol (e.g., Fig. 3 and 4).

## Acknowledgments

The authors thank the patient for participation and Christopher R. Holdgraf, Vinitha Rangarajan, Colin W. Hoy, Julia Kam, Ludovic Bellier, Randolph Helfrich, Richard Jimenez, Edden Gerber, Alejandro Blenkmann, Jamie Lubell, and Michael Pereira for fruitful discussions. The authors are also grateful to the present and former FieldTrip core developers as well as the greater FieldTrip community for contributing code, documentation and expertise that have made this protocol possible. A.S. was supported by Rubicon grant #446-14-007 from NWO and Marie Sklodowska-Curie Global Fellowship #658868 from the European Union; R.vd.M. by R01 #MH095984-03S1 from NIMH; J-M.S. by VIDI #864-14-011 from NWO, R.T.K. by NINDS R37NS21135, and R.O. by H2020-MSCA-ITN-2014 from the EC Marie Curie Actions.

## Author contributions

A.S., S.M.G, R.v.d.M., J-M.S., and R.O. developed the protocol. G.P. contributed the algorithm for brain shift compensation. J.J.L. provided access and guidance in the data acquisition. A.S., S.M.G, J-M.S., R.T.K, and R.O. wrote the paper, and all other authors provided substantial editorial revisions.

---

**TABLE 1.**
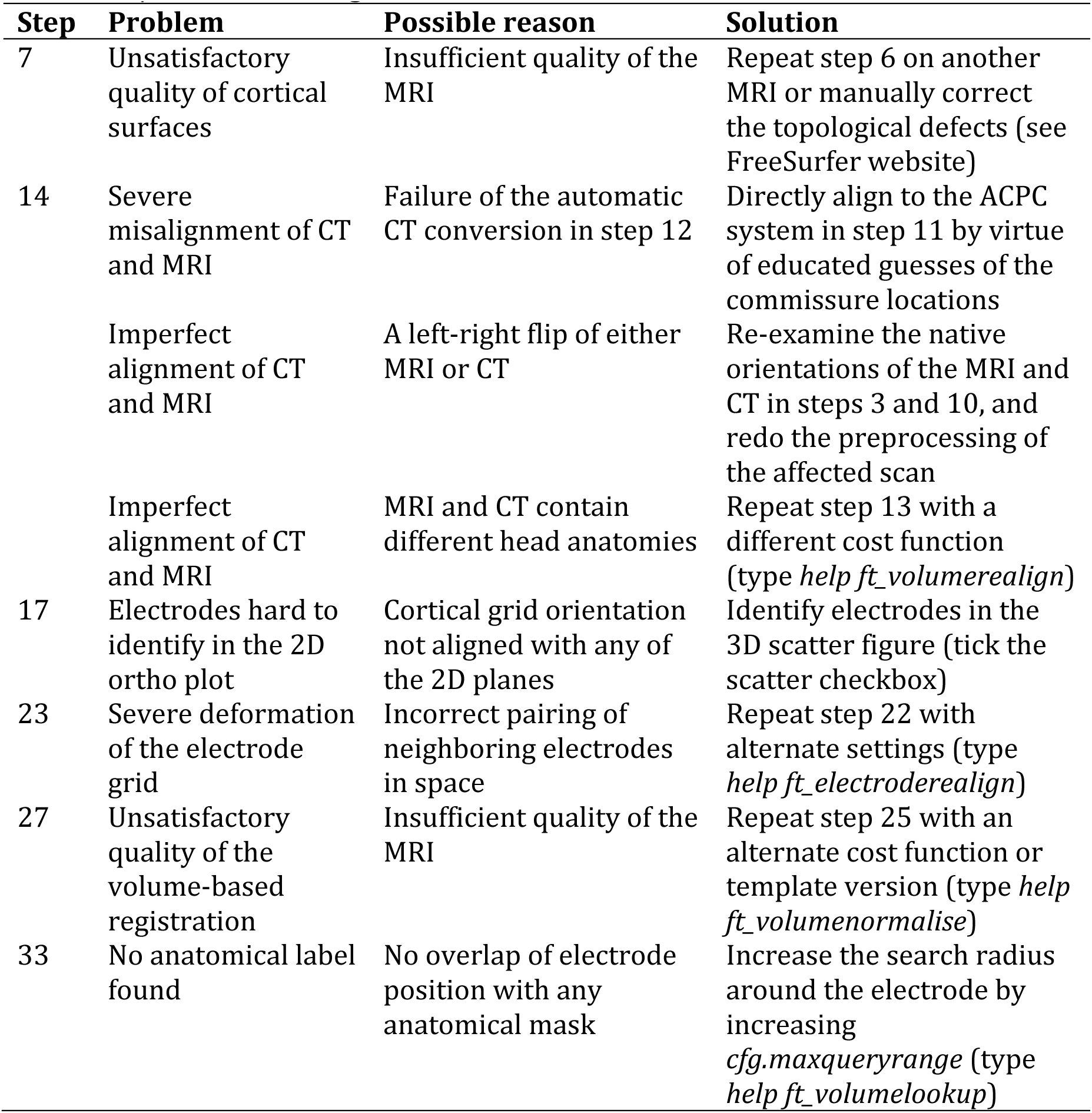
Troubleshooting table.

### Box 1 | Getting started with FieldTrip

All code of the protocol is directly integrated with, and freely available through FieldTrip^53^. This MATLAB-based open source toolbox offers advanced analysis methods for electrophysiological data, such as event-related averaging, frequency and time-frequency analysis, source modeling (for EEG and MEG), connectivity analysis, classification, real-time data processing, and (non)parametric statistical inference. The implementation as a toolbox allows users to perform elaborate and structured analyses of large data sets using the MATLAB command line and batch scripting. Tutorial documentation, answers to frequently asked questions, and example code are available online as a wiki: http://www.fieldtriptoolbox.org. The toolbox’s infrastructure allows users and developers to relatively easily extend the functionality and implement new algorithms. Over the past decade, the FieldTrip toolbox has grown to an estimated 5000 users.

To get started with FieldTrip, download the most recent version from its homepage or GitHub, and set up your MATLAB path.

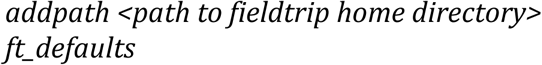

FieldTrip functionalities, recognizable by an *ft* prefix, typically have a single output argument and one or two input arguments, the first input argument being configuration structure *cfg.*

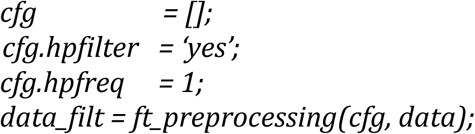

Here, input *data* is processed by *ft_preprocessing* according to parameters specified in the *cfg* fields, in this case applying a 1 Hz high-pass filter. Each function’s optional parameters are available in the respective function’s header (type *help functionname*) and examples are given on the wiki.

The *cfg* structure that holds the parameters to the algorithm at the present level is automatically appended to the output data structure, i.e. *data_filt.cfg.* Configuration structures used at previous levels are kept in *data_filt.cfg.previous*, *data_filt.cfg.previous.previous*, and so on. This nesting of previous configurations allows for access to the full data provenance at any level of the analysis pipeline (see *ft_analysispipeline* for visualizing the pipeline as a flowchart).

### Box 2 | Coordinate system determination, TIMING: ~1 min

Coordinate systems define the orientation and units of the X-, Y-, and Z-axes of an anatomical volume in addition to an origin point along the brain’s midline (e.g., anterior commissure). Here we provide a guideline for determining the native coordinate system of the MRI and CT scans and, in particular, whether they have a left-to-right or a right-to-left orientation. Knowledge of the orientation of the left-right axis of the scan’s native coordinate system provides the necessary context for demarcating the right hemisphere landmark in the succeeding alignment step. Although the interpretation of posterior-anterior and inferior-superior axes is straightforward from axial, coronal, or sagittal slices of the brain, differentiating left and right requires a three-dimensional context. To accomplish this, we recommend using *ft_determine_coordsys,* which depicts an anatomical volume as three intersecting, orthogonal slices and labels the X-, Y-, and Z-axes. This allows determining which of these three axes represents the left-right axis and, importantly, whether that axis has a left-to-right or a right-to-left orientation (Supplementary Video 1).

1. Visualize the coordinate system of the MRI or CT: *ft_determine_coordsys(mri)*
2. Determine which of the three axes, X, Y, or Z, runs through or along the left-right axis of the subject’s head. This axis is the left-right axis for this anatomical volume.
3. Determine the orientation of the left-right axis. If the values on the left-right axis increase to the right (indicated by a + sign), then the scan has a left-to-Right orientation. If the values on the left-right axis increase to the left, then the scan has a right-to-Left orientation.
4. Write down the orientation of the scan’s left-right axis.

### Box 3 | Re-montaging

The recorded electrophysiological signals are a mixture of signal-of-interest and noise, both neural and non-neural. The main objective of the preprocessing of the neural recordings is to improve the signal-to-noise ratio of the data while optimally preparing it for follow-up analysis. Re-montaging to a different referencing scheme, also known as a montage, may aid in the removal of noise that is shared across multiple channels. The common average re-referencing technique, for instance, involves taking the average potential from all channels and subtracting this global noise estimate from the potential in each channel^47, 69–72^. We demonstrated how to apply this technique to the cortical grid electrodes in our example case.

Depth electrodes, located inside the brain and using differently sized and shaped contact points, have a different sensitivity distribution and capture different types of activity and levels of noise^2^. There is currently no consensus on the preferred montage for depth-electrode recordings and, thus, what electrodes to use as references^73–76^. White matter signals may not be as silent as one would intuitively expect, and bipolar signals, despite being relatively clean, miss out on activity that had the same amplitude on the two consecutive electrodes prior to their combination^5, 77^. Different options may need to be tested and evaluated per case, taking into account the purpose of any follow-up analysis^71^. For instance, see ^55^ for a discussion of connectivity analysis in relation to the referencing scheme.

## Supplementary information

Supplementary File **1**. Start-to-end implementation of the anatomical and functional workflows, SubjectUCI29.m

Supplementary File **2**. Automatic DICOM series search and visualization tool, search_dicomseries.m

Supplementary File **3**. Automatic electrode labeling tool, generate_electable.m

Supplementary Video **1**. Preprocessing of the anatomical MRI part 1 https://youtu.be/K9rwGWr-znA

Supplementary Video **2**. Preprocessing of the anatomical MRI part 2 https://youtu.be/KvmiiLY9MiE

Supplementary Video **3**. Preprocessing of the anatomical CT https://youtu.be/or-omFwcswi

Supplementary Video **4**. Electrode placement https://youtu.be/3iaUH7HGUk

Supplementary Video **5**. Interactive plotting https://youtu.be/k7tvP87bnN4

Supplementary Video **6**. Spatiotemporal dynamics of task-modulated high-frequency-band activity at surface electrodes overlaid on left parietal and temporal cortex. It can be observed that processing occurs in the temporal lobe at hearing the target tone followed by the sensorimotor system contralateral to the hand used for the button press. Warm and cold colors represent increases and decreases in high-frequency-band power, respectively. https://youtu.be/PTpive6yiBM

Supplementary Video **7**. Spatiotemporal dynamics of epileptiform activity recorded from depth electrodes targeting bilateral hippocampus and amygdala. It can be observed that the (interictal) epileptiform discharges first occur in the left hippocampus and amygdala and then spread to their right hemisphere homologues during this particular episode. Warm and cold colors represent positive and negative deflections in signal amplitude, respectively. The size of the point clouds indicates the signal’s amplitude. https://youtu.be/7YYumAjLFik

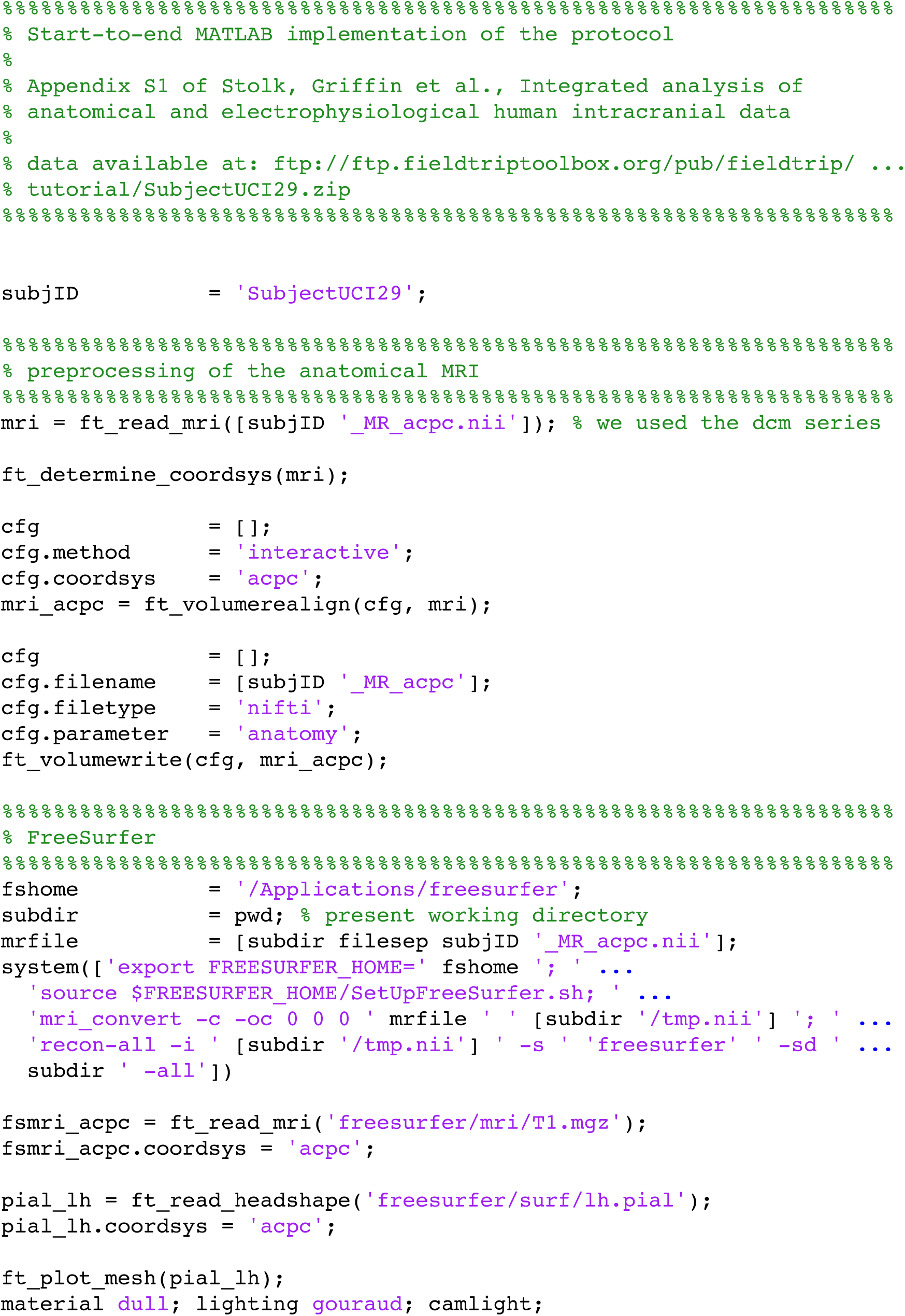

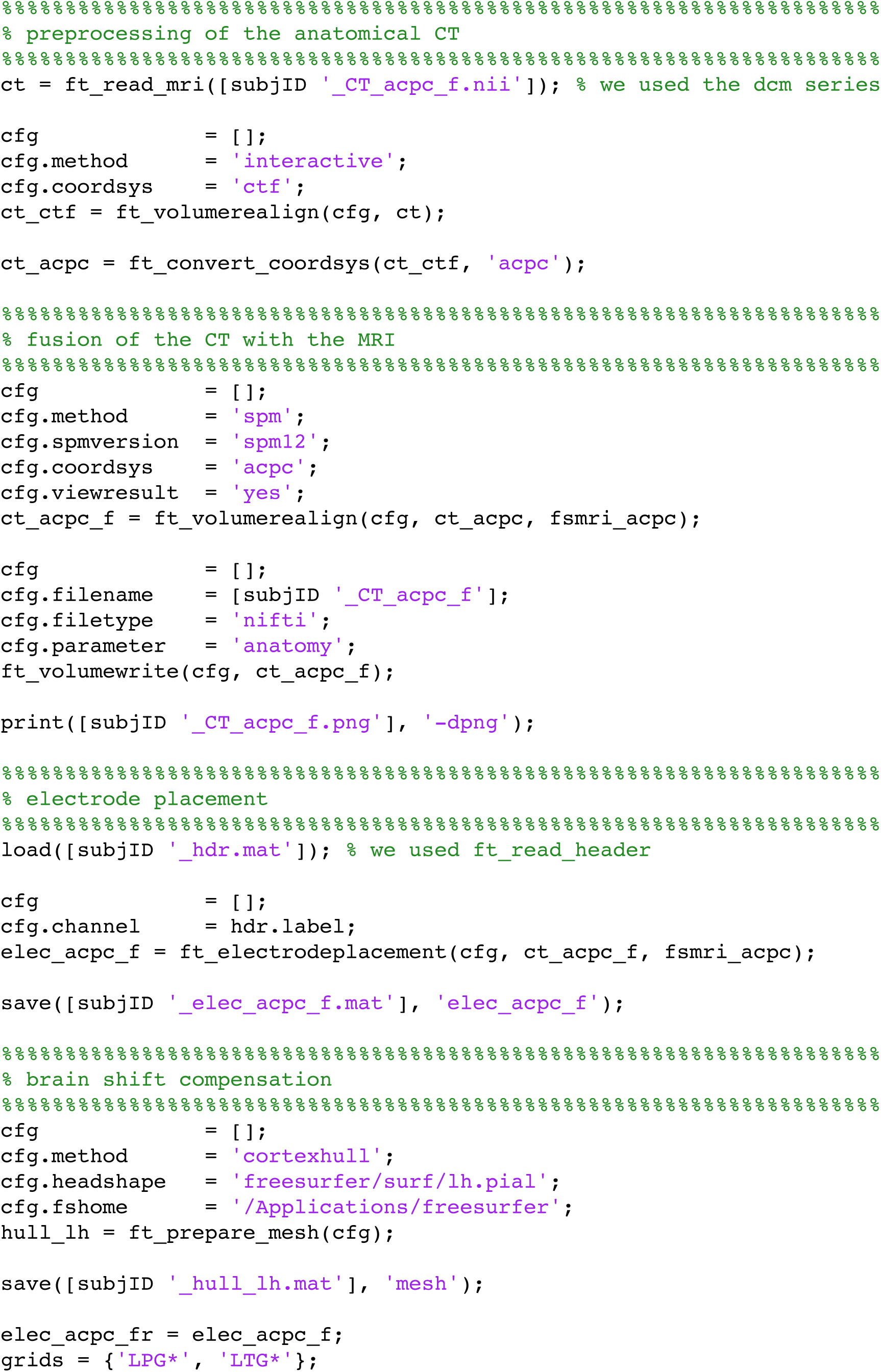

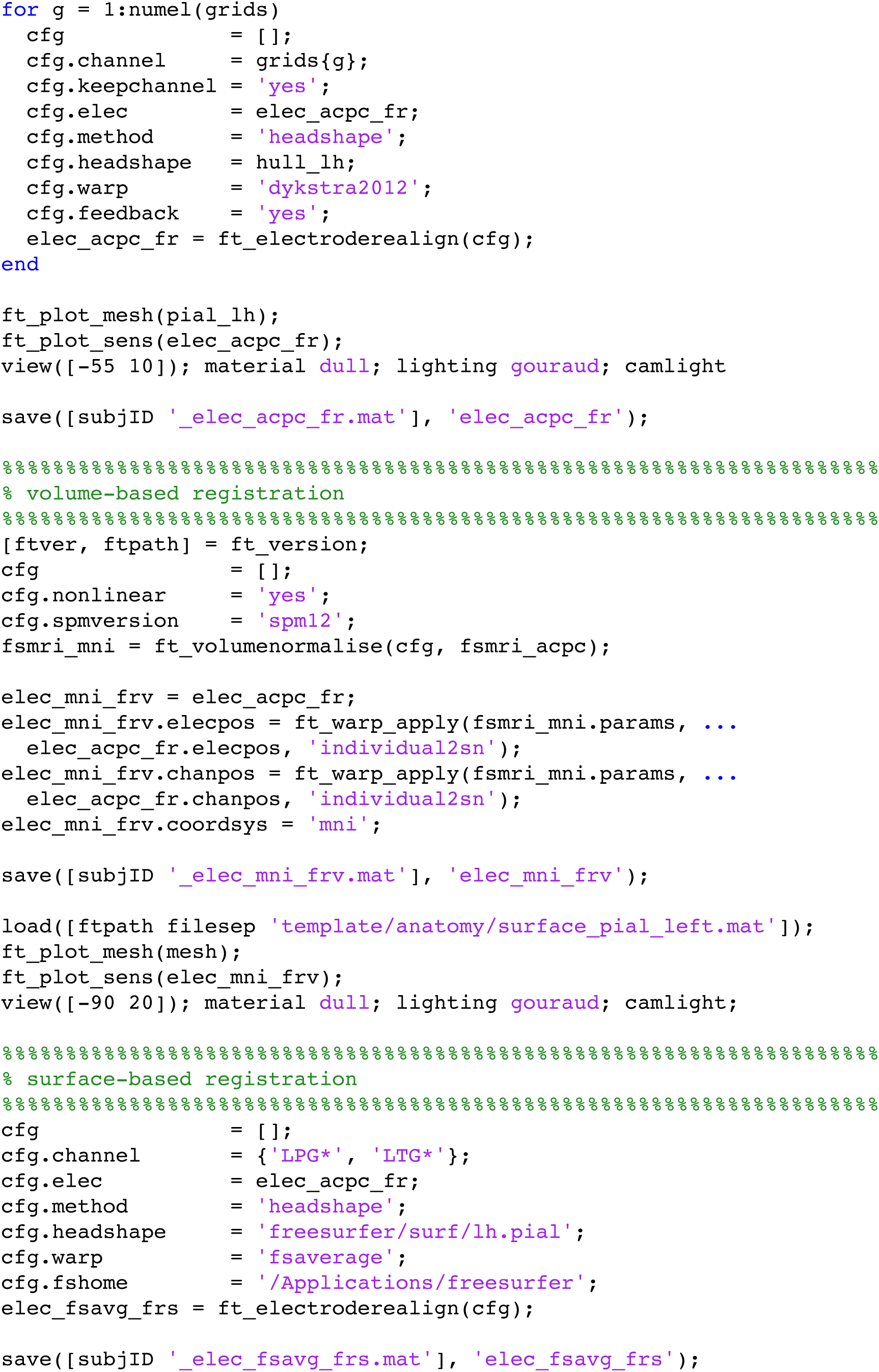

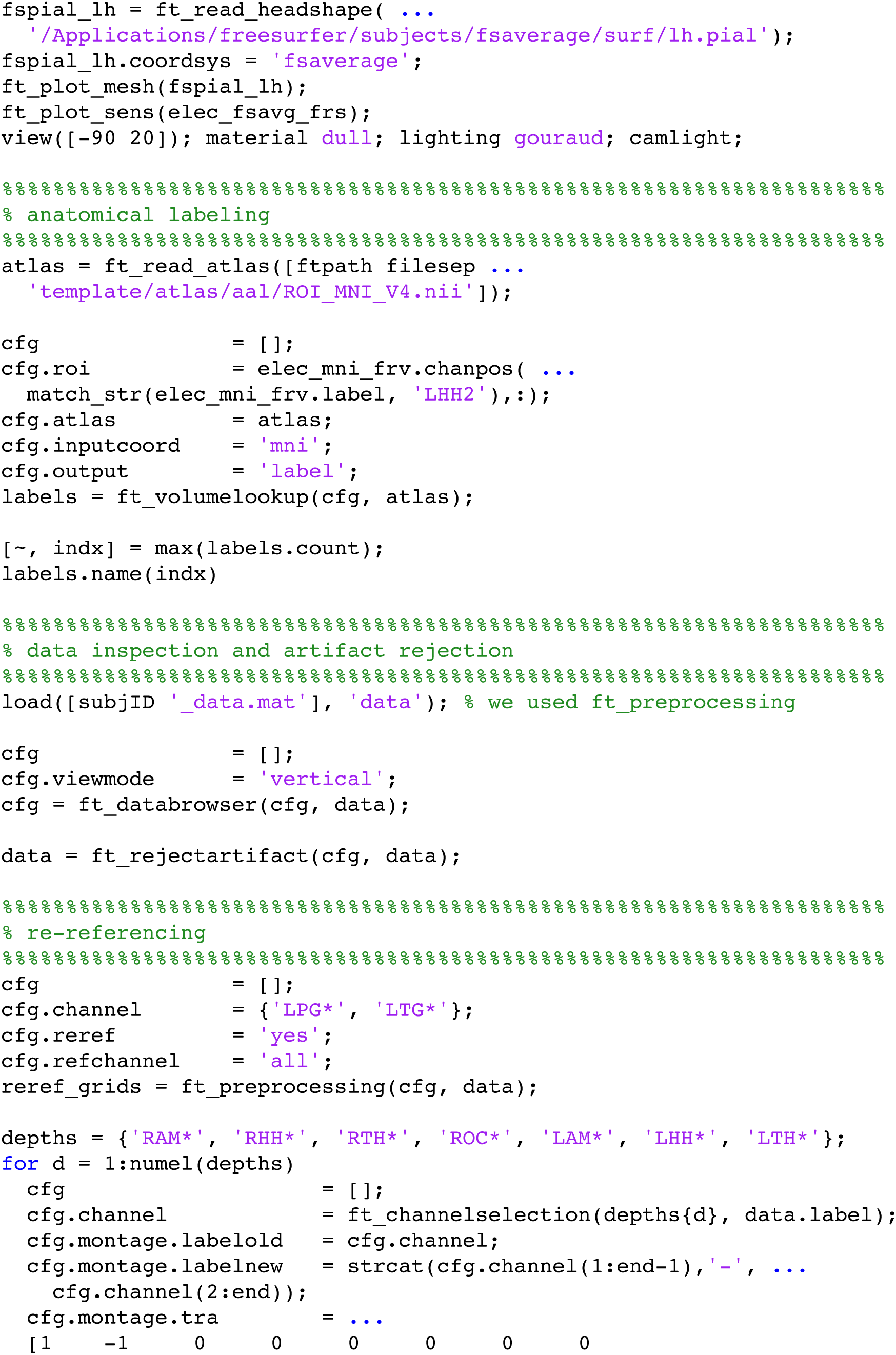

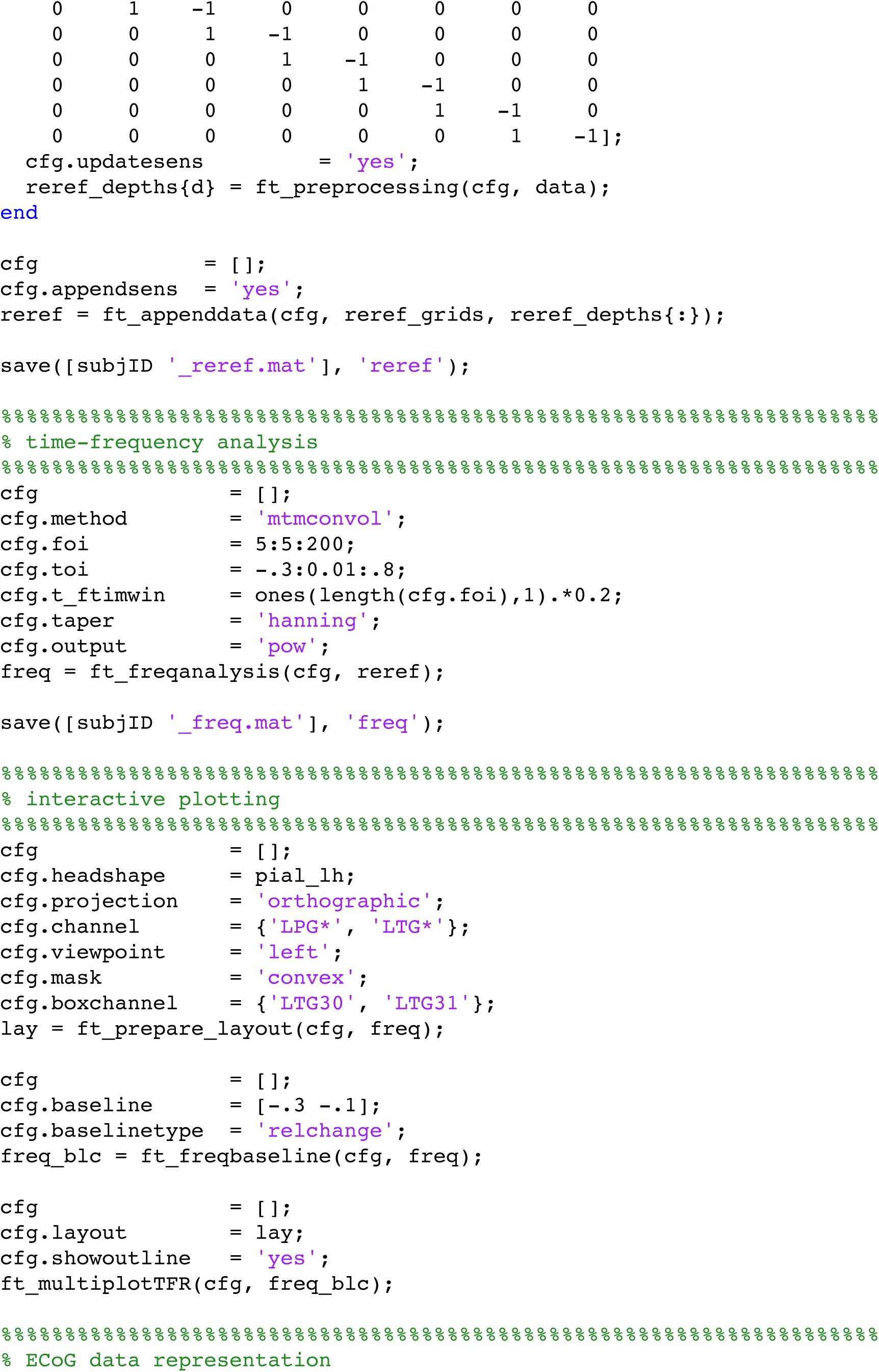

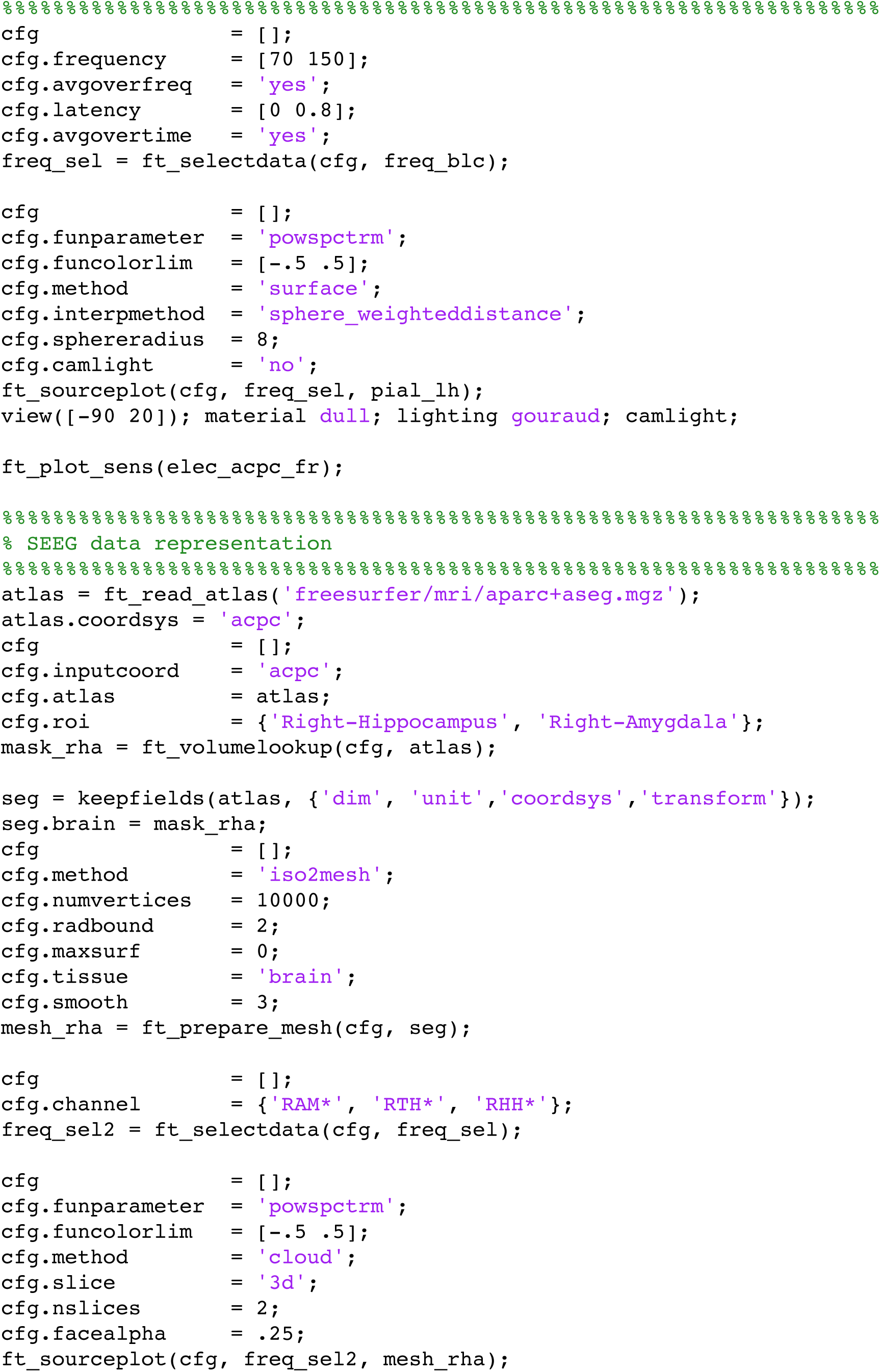

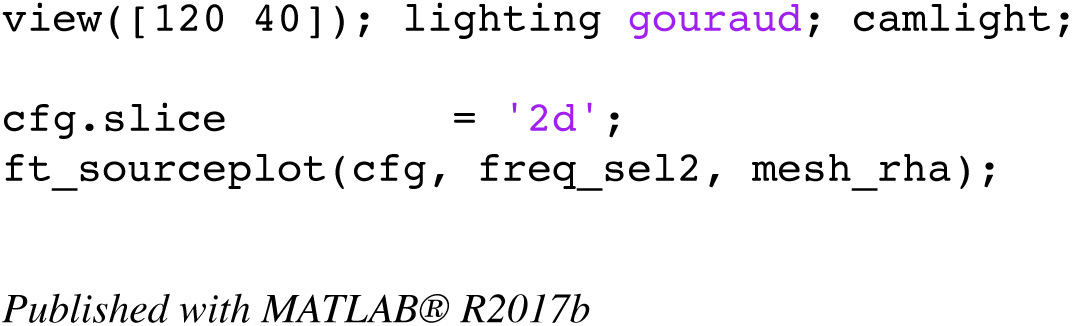

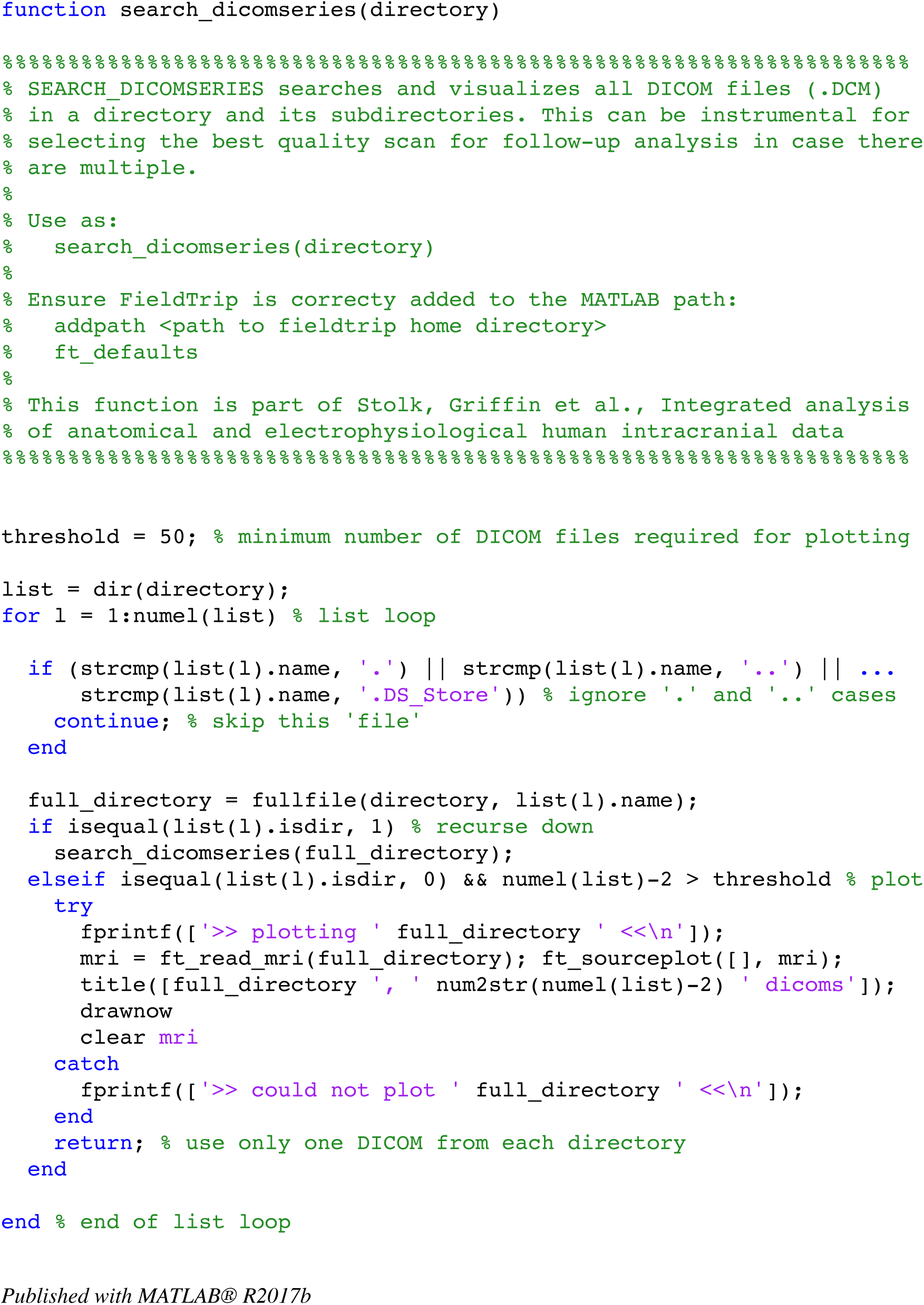

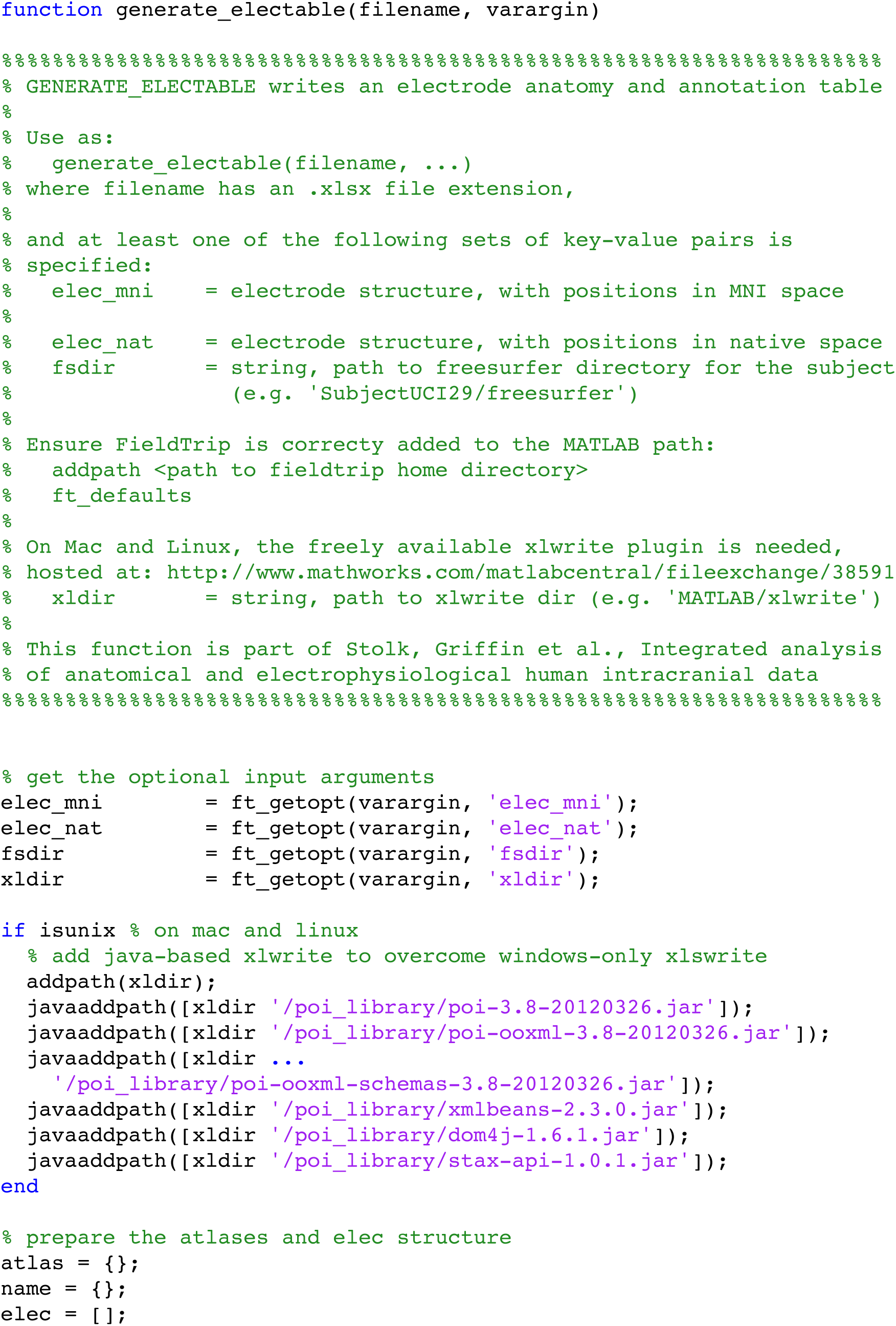

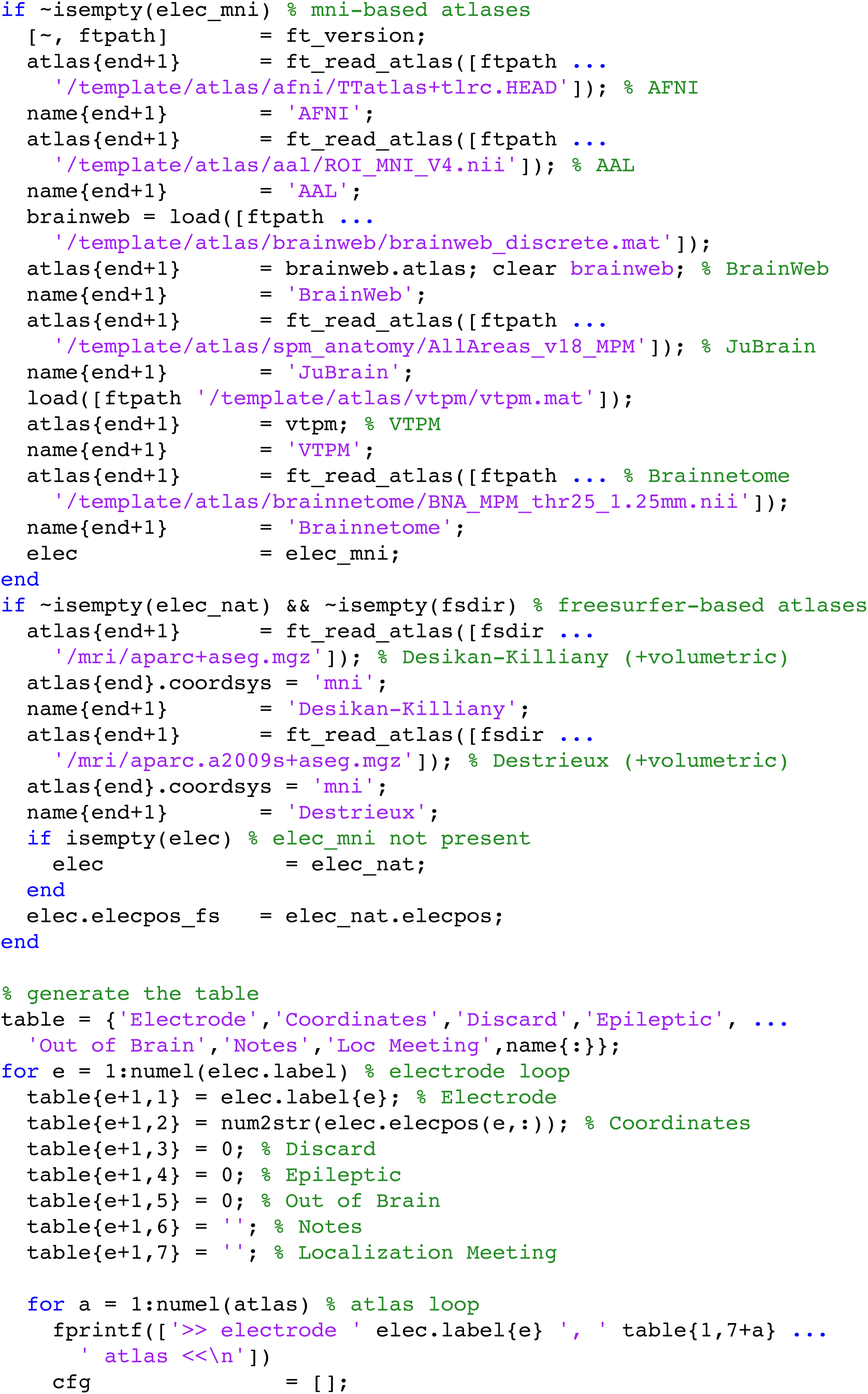

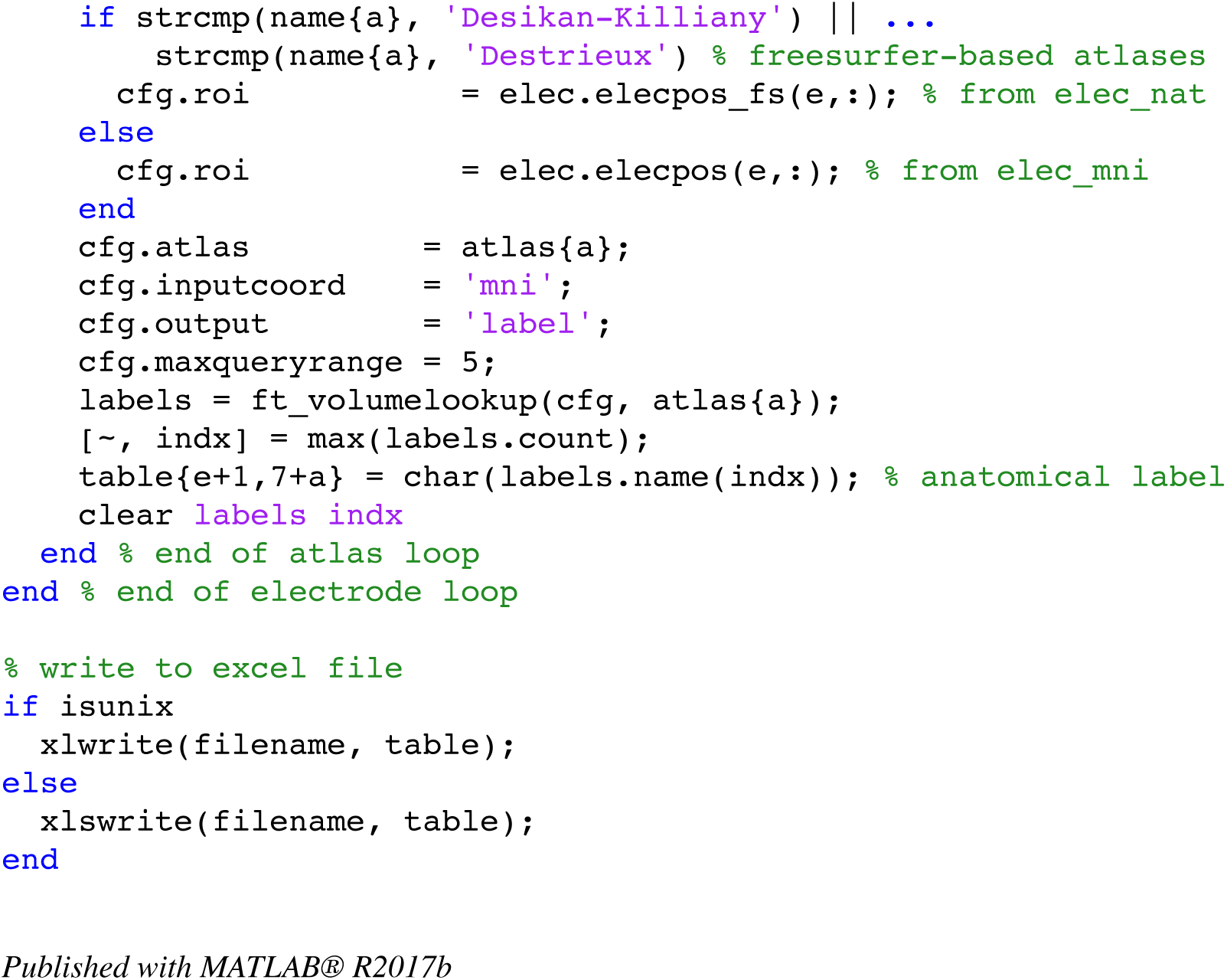

## References

1. Buzsaki, G., Anastassiou, C. A. & Koch, C. The origin of extracellular fields and currents--EEG, ECoG, LFP and spikes. Nat Rev Neurosci 13, 407–420 (2012).

2. Malmivuo, J. & Plonsey, R. Bioelectromagnetism: Principles and Applications of Bioelectric and Biomagnetic Fields. Bioelectromagnetism: Principles and Applications of Bioelectric and Biomagnetic Fields (2012). doi:10.1093/acprof:oso/9780195058239.001.0001

3. Brunner, P. et al. A practical procedure for real-time functional mapping of eloquent cortex using electrocorticographic signals in humans. Epilepsy Behav. 15, 278–286 (2009).

4. Ritaccio, A. et al. Proceedings of the Fifth International Workshop on Advances in Electrocorticography. Epilepsy Behav. E&B; 41, 183–92 (2014).

5. Lachaux, J.-P., Axmacher, N., Mormann, F., Halgren, E. & Crone, N. E. High-frequency neural activity and human cognition: Past, present and possible future of intracranial EEG research. Prog. Neurobiol. 98, 279–301 (2012).

6. Friston, J. A. and K. & Wellcome. Multimodal Image Coregistration and Partitioning - a Unified Framework. Neuroimage 6, 209–217 (1997).

7. Jenkinson, M., Bannister, P., Brady, M. & Smith, S. Improved optimization for the robust and accurate linear registration and motion correction of brain images. Neuroimage 17, 825–841 (2002).

8. Cox, R. W. AFNI: software for analysis and visualization of functional magnetic resonance neuroimages. Comput. Biomed. Res. 29, 162–73 (1996).

9. Papademetris, X. et al. BioImage Suite: An integrated medical image analysis suite: An update. Insight J. 2006, 209 (2006).

10. Azarion, A. A. et al. An open-source automated platform for three-dimensional visualization of subdural electrodes using CT-MRI coregistration. Epilepsia 55, 2028–2037 (2014).

11. Blenkmann, A. O. et al. iElectrodes: A Comprehensive Open-Source Toolbox for Depth and Subdural Grid Electrode Localization. Front. Neuroinform. 11, 14 (2017).

12. Groppe, D. M. et al. iELVis: An open source MATLAB toolbox for localizing and visualizing human intracranial electrode data. J. Neurosci. Methods 281, 40–48 (2017).

13. Kubanek, J. & Schalk, G. NeuralAct: A Tool to Visualize Electrocortical (ECoG) Activity on a Three-Dimensional Model of the Cortex. Neuroinformatics 13, 167–174 (2015).

14. Branco, M. P. et al. ALICE: A tool for Automatic Localization of Intra-Cranial Electrodes for clinical and high-density grids. Prep.

15. Qin, C. et al. Automatic and Precise Localization and Cortical Labeling of Subdural and Depth Intracranial Electrodes. Front. Neuroinform. 11, 1–10 (2017).

16. Hill, N. J. et al. Recording Human Electrocorticographic (ECoG) Signals for Neuroscientific Research and Real-time Functional Cortical Mapping. J. Vis. Exp. (2012). doi:10.3791/3993

17. Eglen, S. J. et al. Toward standard practices for sharing computer code and programs in neuroscience. Nat. Neurosci. 20, 770–773 (2017).

18. Delorme, A. & Makeig, S. EEGLAB: An open source toolbox for analysis of single-trial EEG dynamics including independent component analysis. J. Neurosci. Methods 134, 9–21 (2004).

19. Zheng, J. et al. Amygdala-hippocampal dynamics during salient information processing. Nat. Commun. 8, 14413 (2017).

20. Tang, C., Hamilton, L. S. & Chang, E. F. Intonational speech prosody encoding in the human auditory cortex. Science (80-.). 357, (2017).

21. Martinet, L.-E. et al. Human seizures couple across spatial scales through travelling wave dynamics. Nat. Commun. 8, 14896 (2017).

22. Gelinas, J. N., Khodagholy, D., Thesen, T., Devinsky, O. & Buzsáki, G. Interictal epileptiform discharges induce hippocampal-cortical coupling in temporal lobe epilepsy. Nat. Med. 22, 641–648 (2016).

23. Hermes, D., Miller, K. J., Noordmans, H. J., Vansteensel, M. J. & Ramsey, N. F. Automated electrocorticographic electrode localization on individually rendered brain surfaces. J. Neurosci. Methods 185, 293–298 (2010).

24. Dalal, S. S. et al. Localization of neurosurgically implanted electrodes via photograph-MRI-radiograph coregistration. J. Neurosci. Methods 174, 106–115 (2008).

25. Yang, A. I. et al. Localization of dense intracranial electrode arrays using magnetic resonance imaging. Neuroimage 63, 157–165 (2012).

26. Onofrey, J. A., Staib, L. H. & Papademetris, X. Learning intervention-induced deformations for non-rigid MR-CT registration and electrode localization in epilepsy patients. NeuroImage Clin. 10, 291–301 (2016).

27. Pieters, T. A., Conner, C. R. & Tandon, N. Recursive grid partitioning on a cortical surface model: an optimized technique for the localization of implanted subdural electrodes. J. Neurosurg. 118, 1086–1097 (2013).

28. Stieglitz, L. H. et al. Improved localization of implanted subdural electrode contacts on magnetic resonance imaging with an elastic image fusion algorithm in an invasive electroencephalography recording. Clin. Neurosurg. 10, 506–513 (2014).

29. Brang, D., Dai, Z., Zheng, W. & Towle, V. L. Registering imaged ECoG electrodes to human cortex: A geometry-based technique. J. Neurosci. Methods 273, 64–73 (2016).

30. Dykstra, A. R. et al. Individualized localization and cortical surface-based registration of intracranial electrodes. Neuroimage 59, 3563–3570 (2012).

31. Khodagholy, D. et al. Organic electronics for high-resolution electrocorticography of the human brain. Sci. Adv. 2, 1–9 (2016).

32. Seo, D. et al. Wireless Recording in the Peripheral Nervous System with Ultrasonic Neural Dust. Neuron 91, 529–539 (2016).

33. Lauro, P. M. et al. DBSproc: An open source process for DBS electrode localization and tractographic analysis. Hum. Brain Mapp. 37, 422–433 (2016).

34. Horn, A. & Kühn, A. A. Lead-DBS: A toolbox for deep brain stimulation electrode localizations and visualizations. Neuroimage 107, 127–135 (2015).

35. Dale, A. M., Fischl, B. & Sereno, M. I. Cortical Surface-Based Analysis: I. Segmentation and Surface Reconstruction. Neuroimage 9, 179–194 (1999).

36. Lepore, N. et al. A New Combined Surface and Volume Registration. Med. Imaging 2010 Image Process. 7623, (2010).

37. Klein, A. et al. Evaluation of volume-based and surface-based brain image registration methods. Neuroimage 51, 214–220 (2010).

38. Hill, D. L. G. et al. Measurement of intraoperative brain surface deformation under a craniotomy. Neurosurgery 43, 514–526 (1998).

39. Roberts, D. W., Hartov, A., Kennedy, F. E., Miga, M. I. & Paulsen, K. D. Intraoperative brain shift and deformation: A quantitative analysis of cortical displacement in 28 cases. Neurosurgery 43, 749–758 (1998).

40. Miyagi, Y., Shima, F. & Sasaki, T. Brain shift: an error factor during implantation of deep brain stimulation electrodes. J. Neurosurg. 107, 989–97 (2007).

41. Hastreiter, P. et al. Strategies for brain shift evaluation. Med. Image Anal. 8, 447–464 (2004).

42. LaViolette, P. S. et al. Three-dimensional visualization of subdural electrodes for presurgical planning. Neurosurgery 68, (2011).

43. Sweet, J. A., Hdeib, A. M., Sloan, A. & Miller, J. P. Depths and grids in brain tumors: Implantation strategies, techniques, and complications. Epilepsia 54, 66–71 (2013).

44. Kovalev, D. et al. Rapid and fully automated visualization of subdural electrodes in the presurgical evaluation of epilepsy patients. Am. J. Neuroradiol. 26, 1078–1083 (2005).

45. Wang, P. T. et al. A co-registration approach for electrocorticogram electrode localization using post-implantation MRI and CT of the head. in International IEEE/EMBS Conference on Neural Engineering, NER 525–528 (2013). doi:10.1109/NER.2013.6695987

46. Schulze-Bonhage, A. H. J. et al. Visualization of subdural strip and grid electrodes using curvilinear reformatting of 3D MR imaging data sets. Am. J. Neuroradiol. 23, 400–403 (2002).

47. Boatman-Reich, D. et al. Quantifying auditory event-related responses in multichannel human intracranial recordings. Front. Comput. Neurosci. 4, 4 (2010).

48. Saez, I. et al. Dissociable roles for transient and sustained responses in human OFC during decision-making. Under Rev.

49. Manning, J. R., Jacobs, J., Fried, I. & Kahana, M. J. Broadband shifts in local field potential power spectra are correlated with single-neuron spiking in humans. J Neurosci 29, 13613–13620 (2009).

50. Miller, K. J. Broadband spectral change: evidence for a macroscale correlate of population firing rate? J. Neurosci. 30, 6477–6479 (2010).

51. Ray, S. & Maunsell, J. H. R. Different origins of gamma rhythm and high-gamma activity in macaque visual cortex. PLoS Biol. 9, (2011).

52. Crone, N. E., Miglioretti, D. L., Gordon, B., Lesser, R. P. & Crone, N. Functional mapping of human sensorimotor cortex with electrocorticographic spectral analysis II. Event-related synchronization in the gamma band. Brain 121, 2301–2315 (1998).

53. Oostenveld, R., Fries, P., Maris, E. & Schoffelen, J. M. FieldTrip: Open source software for advanced analysis of MEG, EEG, and invasive electrophysiological data. Comput. Intell. Neurosci. 2011, (2011).

54. Maris, E. & Oostenveld, R. Nonparametric statistical testing of EEG- and MEG-data. J. Neurosci. Methods 164, 177–190 (2007).

55. Bastos, A. M. & Schoffelen, J.-M. A Tutorial Review of Functional Connectivity Analysis Methods and Their Interpretational Pitfalls. Front. Syst. Neurosci. 9, 1–23 (2016).

56. Drury, H. A., Van Essen, D. C., Corbetta, M. & Snyder, A. Z. in Brain Warping 337–363 (1999).

57. Wells, W. M., Viola, P., Atsumi, H., Nakajima, S. & Kikinis, R. Multi-modal volume registration by maximization of mutual information. Med. Image Anal. 1, 35–51 (1996).

58. Collignon, A. & Maes, F. Automated multi-modality image registration based on information theory. Proc. Inf. Process. Med. Imaging 263–274 (1995).

59. Schaer, M. et al. A Surface-based approach to quantify local cortical gyrification. IEEE Trans. Med. Imaging 27, 161–170 (2008).

60. Ashburner, J. & Friston, K. J. Nonlinear spatial normalization using basis functions. Hum. Brain Mapp. 7, 254–266 (1999).

61. Lancaster, J. L. et al. Automated labeling of the human brain: A preliminary report on the development and evaluation of a forward-transform method. in Human Brain Mapping 5, 238–242 (1997).

62. Tzourio-Mazoyer, N. et al. Automated anatomical labeling of activations in SPM using a macroscopic anatomical parcellation of the MNI MRI single-subject brain. Neuroimage 15, 273–289 (2002).

63. Cocosco, C. a, Kollokian, V., Kwan, R. K., Pike, G. B. & Evans, A. C. BrainWeb: Online Interface to a 3D MRI Simulated Brain Database. 3-rd Int. Conf. Funct. Mapp. Hum. Brain 1131, 1996 (1996).

64. Eickhoff, S. B. et al. A new SPM toolbox for combining probabilistic cytoarchitectonic maps and functional imaging data. Neuroimage 25, 1325–1335 (2005).

65. Wang, L., Mruczek, R. E. B., Arcaro, M. J. & Kastner, S. Probabilistic maps of visual topography in human cortex. Cereb. Cortex 25, 3911–3931 (2015).

66. Fan, L. et al. The Human Brainnetome Atlas: A New Brain Atlas Based on Connectional Architecture. Cereb. Cortex 26, 3508–3526 (2016).

67. Desikan, R. S. et al. An automated labeling system for subdividing the human cerebral cortex on MRI scans into gyral based regions of interest. Neuroimage 31, 968–980 (2006).

68. Destrieux, C., Fischl, B., Dale, A. & Halgren, E. Automatic parcellation of human cortical gyri and sulci using standard anatomical nomenclature. Neuroimage 53, 1–15 (2010) .

69. Bigdely-Shamlo, N., Mullen, T., Kothe, C., Su, K.-M. & Robbins, K. A. The PREP pipeline: standardized preprocessing for large-scale EEG analysis. Front. Neuroinform. 9, 16 (2015).

70. Liu, Y., Coon, W. G., Pesters, A. de, Brunner, P. & Schalk, G. The effects of spatial filtering and artifacts on electrocorticographic signals. J. Neural Eng. 12, 56008 (2015).

71. Dien, J. Issues in the application of the average reference: Review, critiques, and recommendations. Behav. Res. Methods, Instruments, Comput. 30, 34–43 (1998).

72. Ludwig, K. A. et al. Using a common average reference to improve cortical neuron recordings from microelectrode arrays. J. Neurophysiol. 101, 1679–1689 (2009).

73. Trongnetrpunya, A. et al. Assessing Granger Causality in Electrophysiological Data: Removing the Adverse Effects of Common Signals via Bipolar Derivations. Front. Syst. Neurosci. 9, 189 (2015).

74. Shirhatti, V., Borthakur, A. & Ray, S. Effect of Reference Scheme on Power and Phase of the Local Field Potential. Neural Comput. 882–913 (2016). doi:10.1162/NECO

75. Arnulfo, G., Hirvonen, J., Nobili, L., Palva, S. & Palva, J. M. Phase and amplitude correlations in resting-state activity in human stereotactical EEG recordings. Neuroimage 112, 114–127 (2015).

76. Zaveri, H. P., Duckrow, R. B. & Spencer, S. S. On the use of bipolar montages for time-series analysis of intracranial electroencephalograms. Clin. Neurophysiol. 117, 2102–2108 (2006).

77. Mercier, M. R. et al. Evaluation of cortical Local Field Potential diffusion in Stereotactic Electro-EncephaloGraphy recordings:a glimpse on white matter signal. Neuroimage (2016). doi:10.1016/j.neuroimage.2016.08.037

